# Antibody structure prediction using interpretable deep learning

**DOI:** 10.1101/2021.05.27.445982

**Authors:** Jeffrey A. Ruffolo, Jeremias Sulam, Jeffrey J. Gray

## Abstract

Therapeutic antibodies make up a rapidly growing segment of the biologics market. However, rational design of antibodies is hindered by reliance on experimental methods for determining antibody structures. In recent years, deep learning methods have driven significant advances in general protein structure prediction. Here, we present DeepAb, a deep learning method for predicting accurate antibody F_V_ structures from sequence. We evaluate DeepAb on two benchmark sets – one balanced for structural diversity and the other composed of clinical-stage therapeutic antibodies – and find that our method consistently outperforms the leading alternatives. Previous deep learning methods have operated as “black boxes” and offered few insights into their predictions. By introducing a directly interpretable attention mechanism, we show that our network attends to physically important residue pairs. For example, in prediction of one CDR H3 residue conformation, the network attends to proximal aromatics and a key hydrogen bonding interaction that constrain the loop conformation. Finally, we present a novel mutant scoring metric derived from network confidence and show that for a particular antibody, all eight of the top-ranked mutations improve binding affinity. These results suggest that this model will be useful for a broad range of antibody prediction and design tasks.

**Significance:** Accurate structure models are critical for understanding the properties of potential therapeutic antibodies. Conventional methods for protein structure determination require significant investments of time and resources and may fail. Although greatly improved, methods for general protein structure prediction still cannot consistently provide the accuracy necessary to understand or design antibodies. We present a deep learning method for antibody structure prediction and demonstrate improvement over alternatives on diverse, therapeutically relevant benchmarks. In addition to its improved accuracy, our method reveals interpretable outputs about specific amino acids and residue interactions that should facilitate design of novel therapeutic antibodies.

## Introduction

The adaptive immune system of vertebrates is capable of mounting robust responses to a broad range of potential pathogens. Critical to this flexibility are antibodies, which are specialized to recognize a diverse set of molecular patterns with high affinity and specificity. This natural role in the defense against foreign particles makes antibodies an increasingly popular choice for therapeutic development^1,2^. Presently, the design of therapeutic antibodies comes with significant barriers^1^. For example, the rational design of antibody-antigen interactions often depends upon an accurate model of antibody structure. However, experimental methods for protein structure determination such as crystallography, NMR, and cryo-EM are low-throughput and time consuming.

Antibody structure consists of two heavy and two light chains that assemble into a large Y-shaped complex. The crystallizable fragment (F_C_) region is involved in immune effector function and is highly conserved within isotypes. The variable fragment (F_V_) region is responsible for antigen binding through a set of six hypervariable loops that form a complementarity determining region (CDR). Structural modeling of the F_V_ is critical for understanding the mechanism of antigen binding and for rational engineering of specific antibodies. Most methods for antibody F_V_ structure prediction employ some form of grafting, by which pieces of previously solved F_V_ structures with similar sequences are combined to form a predicted model^3–6^. Because much of the F_V_ is structurally conserved, these techniques are typically able to produce models with an overall root mean squared deviation (RMSD) less than 1 Å from the native structure. However, the length and conformational diversity of the third CDR loop of the heavy chain (CDR H3) make it difficult to identify high-quality templates. Further, the H3 loop’s position between the heavy and light chains makes it dependent on the chain orientation and multiple adjacent loops^7,8^. Thus the CDR H3 loop presents a longstanding challenge for F_V_ structure prediction methods^9^.

Machine learning methods have become increasing popular for protein structure prediction and design problems^10^. Specific to antibodies^11^, machine learning has been applied to predict developability^12^, improve humanization^13^, generate sequence libraries^14^, and predict antigen interactions^15,16^. In this work, we build on advances in general protein structure prediction^17–19^ to predict antibody F_V_ structures. Our method consists of a deep neural network for predicting inter-residue distances and orientations and a Rosetta-based protocol for generating structures from network predictions. We show that deep learning approaches can predict more accurate structures than grafting-based alternatives, particularly for the challenging CDR H3 loop. The network used in this work is designed to be directly interpretable, providing insights that could assist in structural understanding or antibody engineering efforts. We conclude by demonstrating the ability of our network to distinguish mutational variants with improved binding using a prediction confidence metric. To facilitate further studies, all the code for this work, as well as pre-trained models, are provided.

## Results

### Overview of the method

Our method for antibody structure prediction, DeepAb, consists of two main stages (Figure 1). The first stage is a deep residual convolutional network that predicts F_V_ structure, represented as relative distances and orientations between pairs of residues. The network requires only heavy and light chain sequences as input and is designed with interpretable components to provide insight into model predictions. The second stage is a fast Rosetta-based protocol for structure realization using the predictions from the network.

**Figure 1.**
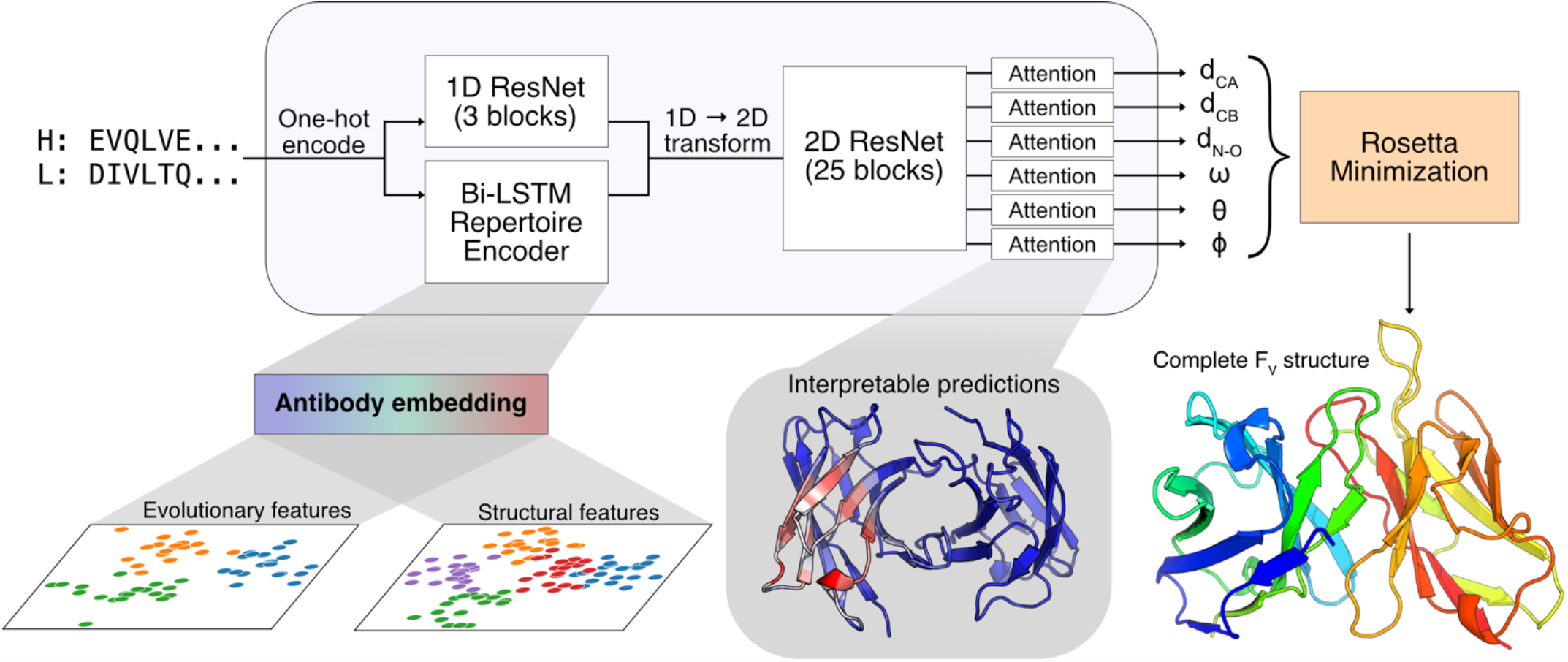
Diagram of DeepAb method for antibody structure prediction. Starting from heavy and light chain sequences, the network predicts a set of inter-residue geometries describing the F_V_ structure. Predictions are used for guided structure realization with Rosetta. Two interpretable components of the network are highlighted: a pre-trained antibody sequence model and output attention mechanisms.

#### Predicting inter-residue geometries from sequence

Due to the limited number of F_V_ crystal structures available for supervised learning, we sought to make use of the abundant immunoglobin sequences from repertoire sequencing studies^20^. We leveraged the power of unsupervised representation learning to extract general patterns from immunoglobin sequences that are not evident in the small subset with known structures. Although transformer models have become increasingly popular for unsupervised learning on protein sequences^21–24^, we chose a recurrent neural network (RNN) model for its relative simplicity and ease of training. The fixed-size hidden state of RNNs forms an explicit information bottleneck ideal for representation learning. In the recent UniRep method, RNNs were demonstrated to learn rich feature representations from protein sequences when trained on next-amino-acid prediction^25^. For our purposes, we developed an RNN encoder-decoder model^26^; the encoder is a bi-LSTM and the decoder is an LSTM^27^. Briefly, the encoder learns to summarize an input sequence residue-by-residue into a fixed-size hidden state. This hidden state is transformed into a summary vector and passed to the decoder, which learns to reconstruct the original sequence one residue at a time. The model is trained using cross-entropy loss on a set of 118,386 paired heavy and light chain sequences from the Observed Antibody Space (OAS) database^28^. After training the network, we generated embeddings for antibody sequences by concatenating the encoder hidden states for each residue. These embeddings are used as features for the structure prediction model described below.

The choice of protein structure representation is critical for structure prediction methods^10^. We represent the F_V_ structure as a set of inter-residue distances and orientations, similar to previous methods for general protein structure prediction^18,19^. Specifically, we predict inter-residue distances between three pairs of atoms (C_α_—C_α,_ C_β_—C_β_, N—O) and the set of inter-residue dihedral and planar angles (*ω, θ, ϕ*) first described by Yang et al^18^. Each output geometry is discretized into 36 bins, with an additional bin indicating distant residue pairs 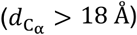. All distances are predicted in the range of [0-18 Å], with a bin width of 0.5 Å. Dihedral and planar angles are discretized uniformly into bins of 10° and 5°, respectively.

The general architecture of the structure prediction network is similar to our previous method for CDR H3 loop structure prediction^29^, with two notable additions: embeddings from the pre-trained language model and interpretable attention layers (Figure 1). The network takes as input the concatenated heavy and light chain sequences. The concatenated sequence is one-hot encoded and passed through two parallel branches: a 1D ResNet and the pre-trained language model. The outputs of the branches are combined and transformed into pairwise data. The pairwise data pass through a deep 2D ResNet that constitutes the main component of the predictive network. Following the 2D ResNet, the network separates into six output branches, corresponding to each type of geometric measurement. Each output branch includes a recurrent criss-cross attention module, allowing each residue pair in the output to aggregate information from all other residue pairs. The attention layers provide interpretability that is often missing from protein structure prediction models.

We opted to train with focal loss^30^ rather than cross-entropy loss to improve the calibration of model predictions, as models trained with cross-entropy loss have been demonstrated to overestimate the likelihood of their predicted labels^31^. Model calibration may be of limited importance for the structure prediction task. However, later in this work we attempt to distinguish between potential antibody variants on the basis of prediction confidence, which requires greater calibration. The model is trained on a nonredundant (at 99% sequence identity) set of 1,692 F_V_ structures from the Structural Antibody Database (SAbDab)^32^. The pretrained language model, used as a feature extractor, is not updated while training the predictor network.

#### Structure realization

Similar to previous methods for general protein structure prediction^17–19^, we used constrained minimization to generate full 3D structures from network predictions. Unlike previous methods, which typically begin with some form of *ϕ* − *ψ* torsion sampling, we created initial models via multi-dimensional scaling (MDS). As a reminder, the relative positions of all backbone atoms are fully specified by the predicted *L* × *L* inter-residue 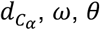, and *ϕ* geometries. Using the modal-predicted output bins for these four geometries, we construct a distance matrix between backbone atoms. From this distance matrix, MDS produces an initial set of 3D coordinates that are subsequently refined through constrained minimization.

Network predictions for each output geometry were converted to energetic potentials by negating the raw model logits (i.e., without softmax activation). These discrete potentials were converted to continuous constraints using a cubic spline function. Starting from the MDS model, the constraints are used to guide quasi-Newton minimization (L-BFGS) within Rosetta^33,34^. First, the constraints are jointly optimized with the Rosetta centroid energy function to produce a coarse-grained F_V_ structure with the sidechains represented as a single atom. Next, constrained full-atom relaxation was used to introduce sidechains and remove clashes. After relaxation, the structure was minimized again with constraints and the Rosetta full-atom energy function (ref2015). This optimization procedure was repeated to produce 50 structures, and the lowest-energy structure was selected as the final model. Although we opted to produce 50 candidate structures, 5-10 should be sufficient in practice due to the high convergence of the protocol (Figure S1).

### Benchmarking methods for F_V_ structure prediction

To evaluate the performance of our method, we chose two independent test sets. The first is the RosettaAntibody benchmark set, which has previously been used to evaluate antibody structure prediction methods^8,29,35^. The second is a set of clinical-stage therapeutic antibodies, which was previously assembled to study antibody developability^36^. Taken together, these sets represent a structurally diverse, therapeutically relevant benchmark for comparing antibody F_V_ structure prediction methods.

#### Deep learning outperforms grafting methods

We compared the performance of our method on the RosettaAntibody benchmark and therapeutic benchmark to three alternative methods: RosettaAntibody-G^4,6^, RepertoireBuilder^5^, and ABodyBuilder^3^. Each of these methods is based on a grafting approach, by which complete F_V_ structures are assembled from sequence-similar fragments of previously solved structures. To produce the fairest comparison, we excluded structures with greater than 99% sequence identity for the whole F_V_ from use for grafting (similar to our training dataset). We evaluated each method according to the backbone heavy-atom RMSD of the CDR loops and the framework regions of both chains. We also measured the orientational coordinate distance (OCD)^8^, a metric for heavy-light chain orientation accuracy. OCD is calculated as the sum of the deviations from native of four orientation coordinates (packing angle, interdomain distance, heavy-opening angle, light-opening angle) divided by the standard deviation of each coordinate^8^. The results of the benchmark are summarized in Table 1 and fully detailed in Tables S1-8.

**Table 1.** Performance of F_V_ structure prediction methods on benchmarks.

| Method | OCD | H Fr (Å) | H1 (Å) | H2 (Å) | H3 (Å) | L Fr (Å) | L1 (Å) | L2 (Å) | L3 (Å) |
| --- | --- | --- | --- | --- | --- | --- | --- | --- | --- |
| RosettaAntibody Benchmark |  |  |  |  |  |  |  |  |  |
| RosettaAntibody-G | 5.19 | 0.57 | 1.22 | 1.14 | 3.48 | 0.67 | 0.80 | 0.87 | 1.06 |
| RepertoireBuilder | 5.26 | 0.58 | 0.86 | 1.00 | 2.94 | 0.51 | 0.63 | 0.52 | 1.03 |
| ABodyBuilder | 4.69 | 0.50 | 0.99 | 0.88 | 2.94 | 0.49 | 0.72 | 0.52 | 1.09 |
| DeepAb | **3.67** | **0.43** | **0.72** | **0.85** | **2.33** | **0.42** | **0.55** | **0.45** | **0.86** |
| Therapeutic Benchmark |  |  |  |  |  |  |  |  |  |
| RosettaAntibody-G | 5.43 | 0.63 | 1.42 | 1.05 | 3.77 | 0.55 | 0.89 | 0.83 | 1.48 |
| RepertoireBuilder | 4.37 | 0.62 | 0.91 | 0.96 | 3.13 | 0.47 | 0.71 | 0.52 | 1.08 |
| ABodyBuilder | 4.37 | 0.49 | 1.05 | 1.02 | 3.00 | 0.45 | 1.04 | 0.50 | 1.35 |
| DeepAb | **3.52** | **0.40** | **0.77** | **0.68** | **2.52** | **0.37** | **0.60** | **0.42** | **1.02** |

Orientational coordinate distance (OCD) is unitless quantity calculated by measuring the deviation from native of four heavy-light chain coordinates<sup>1</sup>. Heavy chain framework (H Fr) and light chain framework (L Fr) RMSDs are measured after superimposing the heavy and light chains, respectively. CDR loop RMSDs are measured using the Chothia loop definitions after superimposing the framework region of the corresponding chain. All RMSDs are measured over backbone heavy atoms.

Our deep learning method showed improvement over all grafting-based methods on every metric considered. On both benchmarks, the structures predicted by our method achieved an average OCD less than 4, indicating that predicted structures were typically within one standard deviation of the native structure for each of the orientational coordinates. All of the methods predicted with sub-Angstrom accuracy on the heavy and light chain framework regions, which are highly conserved. Still, our method achieved average RMSD improvements of 14-18% for the heavy chain framework and 16-17% for light chain framework over the next best methods on the benchmarks. We also observed consistent improvement over grafting methods for CDR loop structure prediction.

#### Comparison of CDR H3 loop modeling accuracy

The most significant improvements by our method were observed for the CDR H3 loop (Figure 2A). On the RosettaAntibody benchmark, our method predicted H3 loop structures with an average RMSD of 2.33 Å (± 1.32 Å), a 22% improvement over the next best method. On the therapeutic benchmark, our method had an average H3 loop RMSD of 2.52 Å (± 1.50 Å), a 16% improvement over the next best method. The difficulty of predicting CDR H3 loop structures is due in part to the wide range of observed loop lengths. To understand the impact of H3 loop length on our method’s performance, we compared the average RMSD for each loop length across both benchmarks (Figure 2B). In general, all of the methods displayed degraded performance with increasing H3 loop length. However, DeepAb typically produced the most accurate models for each loop length.

**Figure 2.**
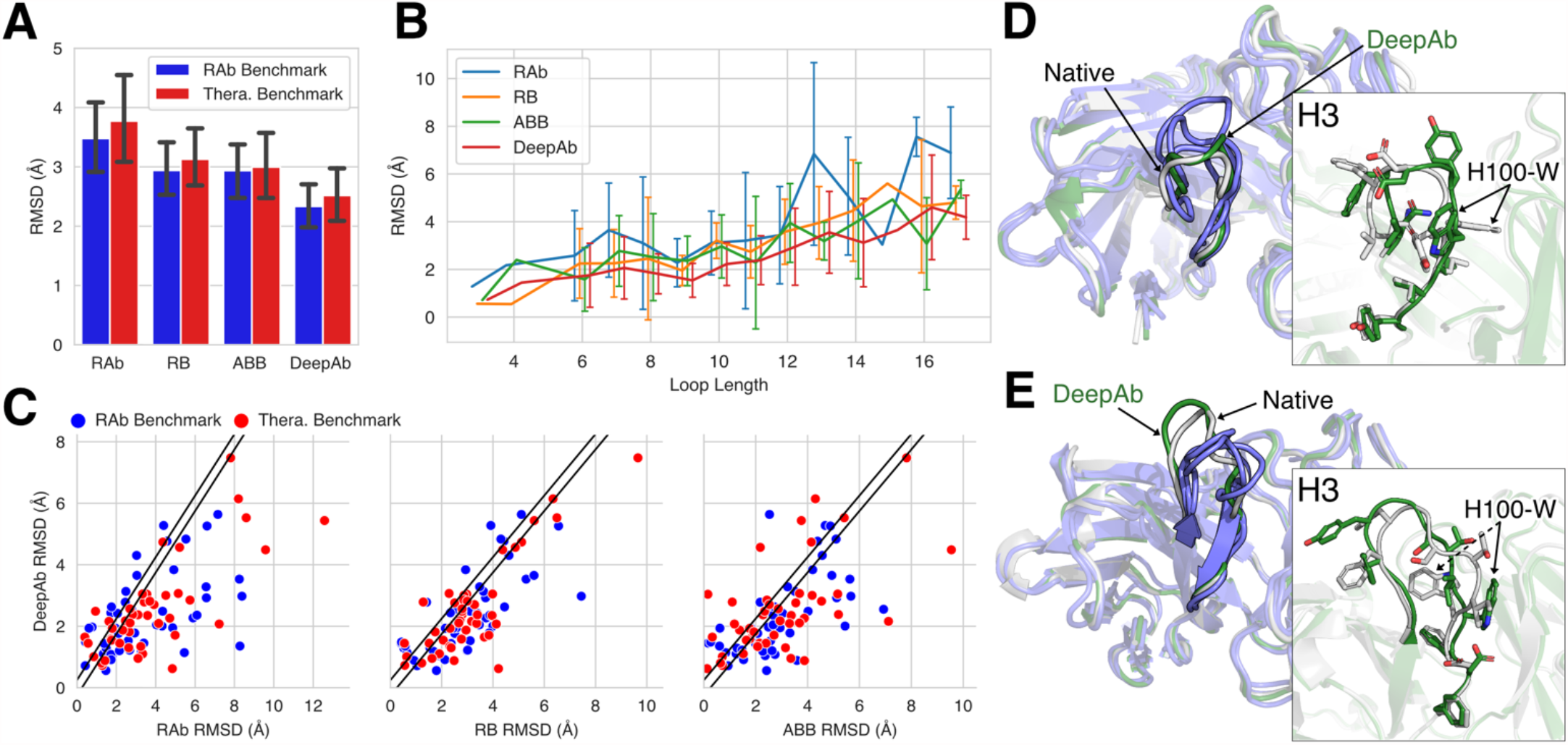
Comparison of CDR H3 loop structure prediction accuracy. (A) Average RMSD of H3 loops predicted by RosettaAntibody-G (RAb), RepertoireBuilder (RB), ABodyBuilder (ABB), and DeepAb on the two benchmarks. Error bars show standard deviations for each method on each benchmark. (B) Average RMSD of H3 loops by length for all benchmark targets. Error bars show standard deviations for loop lengths corresponding to more than one target. (C) Direct comparison of DeepAb and alternative methods H3 loop RMSDs, with diagonal band indicating predictions that were within ±0.25 Å. (D) Comparison of native rituximab H3 loop structure (white, PDB ID 3PP3) to predictions from DeepAb (green, 2.1 Å RMSD) and alternative methods (blue, 3.3-4.1 Å RMSD). (E) Comparison of native sonepcizumab H3 loop structure (white, PDB ID 3I9G) to predictions from DeepAb (green, 1.8 Å RMSD) and alternative methods (blue, 2.9-3.9 Å RMSD).

We also examined the performance of each method on individual benchmark targets. In Figure 2C, we plot the CDR H3 loop RMSD of our method versus that of the alternative methods. Predictions with an RMSD difference less than 0.25 Å (indicated by diagonal bands) were considered equivalent in quality. When compared to RosettaAntibody-G, RepertoireBuilder, and ABodyBuilder, our method predicted more/less accurate H3 loop structures for 64/17, 59/16, and 53/22 out of 92 targets, respectively. Remarkably, our method was able to predict nearly half of the H3 loop structures (42 out of 92) to within 2 Å RMSD. RosettaAntibody-G, RepertoireBuilder, and ABodyBuilder achieved RMSDs of 2 Å or better on 26, 23, and 26 targets, respectively.

#### Accurate prediction of challenging, therapeutically relevant targets

To underscore and illustrate the improvements achieved by our method, we highlight two examples from the benchmark sets. The first is rituximab, an anti-CD20 antibody from the therapeutic benchmark (PDB ID 3PP3)^37^. In Figure 2D, the native structure of the 12-residue rituximab H3 loop (white) is compared to our method’s prediction (green, 2.1 Å RMSD) and the predictions from the grafting methods (blue, 3.3-4.1 Å RMSD). The prediction from our method captures the general topology of the loop well, even placing many of the side chains near the native. The second example is sonepcizumab, an anti-sphingosine-1-phosphate antibody from the RosettaAntibody benchmark (PDB ID 3I9G)^38^. In Figure 2E, the native structure of the 12-residue H3 loop (white) is compared to our method’s prediction (green, 1.8 Å) and the predictions from the grafting methods (blue, 2.9-3.9 Å). Again, our method captures the overall shape of the loop well, enabling accurate placement of several side chains. Interestingly, the primary source of error by our method in both cases is a tryptophan residue (around position H100) facing in the incorrect direction.

### Interpretability of model predictions

Despite the popularity of deep learning approaches for protein structure prediction, little attention has been paid to model interpretability. Interpretable models offer utility beyond their primary predictive task. The network used in this work was designed to be directly interpretable and should be useful for structural understanding and antibody engineering.

#### Output attention tracks model focus

Each output branch in the network includes a criss-cross attention module^39^. The criss-cross attention operation allows the network to attend across output rows and columns when predicting for each residue pair (as illustrated in Figure 3A). Through the attention layer, we create a matrix ***A****∈*ℝ^*L*×*L*^ (where *L* is the total number of residues in the heavy and light chain Fv domains) containing the total attention between each pair of residues (see Methods). To illustrate the interpretative power of network attention, we considered an anti-peptide antibody (PDB ID 4H0H) from the RosettaAntibody benchmark set. Our method performed well on this example (H3 RMSD = 1.2 Å), so we expected it would provide insights into the types of interactions that the network captures well. We collected the attention matrix for 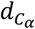 predictions and averaged over the residues belonging to each CDR loop to determine which residues the network focuses on while predicting each loop’s structure (Figure 3B). As expected, the network primarily attends to residues surrounding each loop of interest. For the CDR1-2 loops, the network attends to the residues in the neighborhood of the loop, with little attention paid to the opposite chain. For the CDR3 loops, the network attends more broadly across the heavy-light chain interface, reflecting the interdependence between the loop conformations and the overall orientation of the chains.

**Figure 3.**
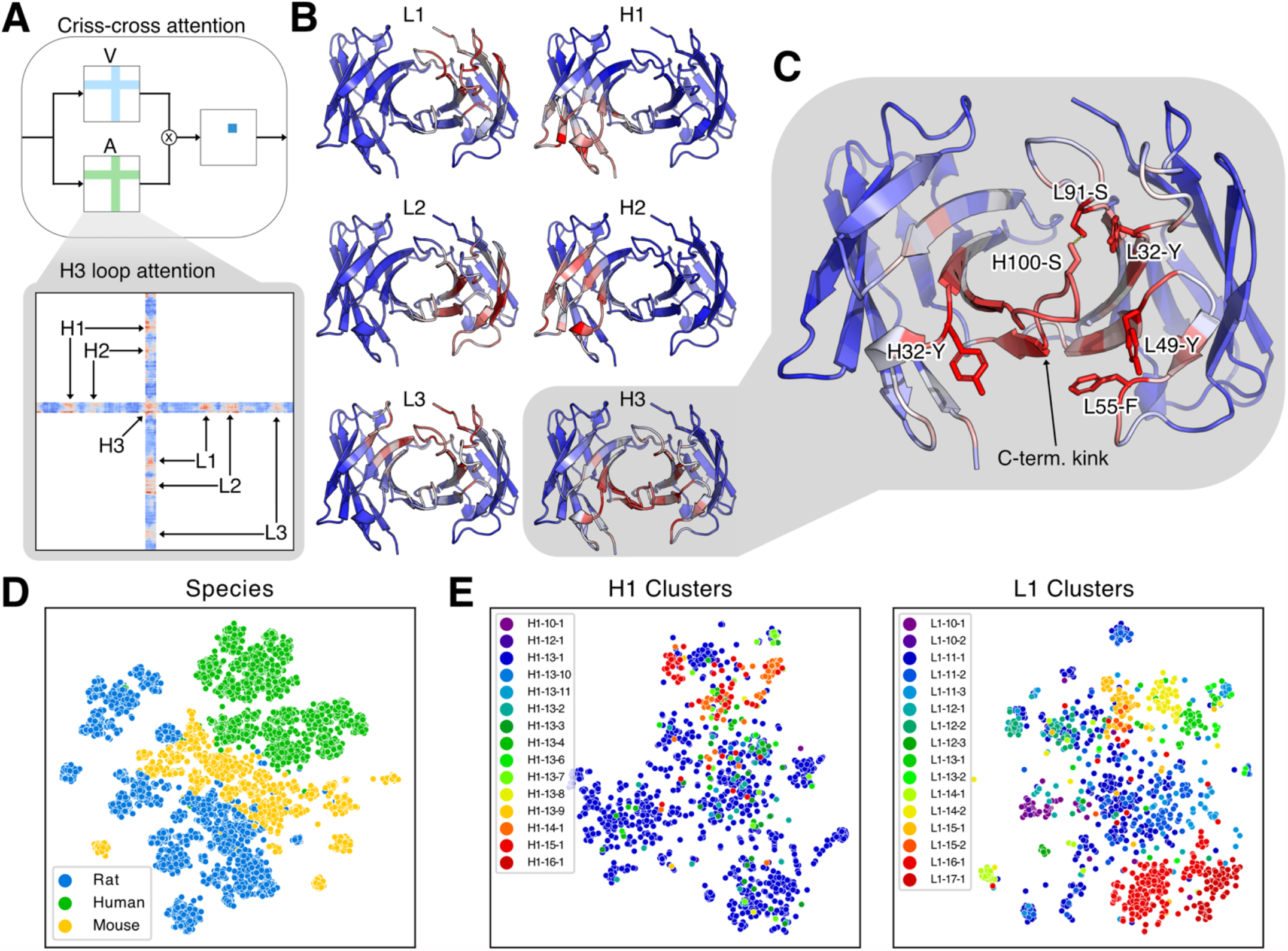
Interpretability of model components. (A) Diagram of attention mechanism (with attention matrix A and value matrix V) and example H3 loop attention matrix, with attention on other loops indicated. Attention values increase from blue to red. (B) Model attention over F_V_ structure while predicting each CDR loop for an anti-peptide antibody (PDB ID 4H0H). (C) Key interactions for H3 loop structure prediction identified by attention. The top five non-H3 attended residues (H32-Y, L32-Y, L49-Y, L55-F, and L91-S) are labeled, as well as an H3 residue participating in a hydrogen bond (H100-S). (D) Two-dimensional t-SNE projection of sequence-averaged LSTM embeddings labeled by source species. (E) Two-dimensional t-SNE projects of LSTM embeddings averaged over CDR1 loop residues labeled by loop structural clusters.

To better understand what types of interactions the network considers, we examined the residues assigned high attention while predicting the H3 loop structure (Figure 3C). Within the H3 loop, we found that the highest attention was on the residues forming the C-terminal kink. This structural feature has previously been hypothesized to contribute to H3 loop conformational diversity^40^, and it is likely critical for correctly predicting the overall loop structure. Of the five non-H3 residues with the highest attention, we found that one was a phenylalanine and three were tyrosines. The coordination of these bulky side chains appears to play a significant role in the predicted H3 loop conformation. The fifth residue was a serine from the L3 loop (residue L91) that forms a hydrogen bond with a serine of the H3 loop (residue H100), suggesting some consideration by the model of biophysical interactions between neighboring residues. To understand how the model attention varies across different H3 loops and neighboring residues, we performed a similar analysis for the 47 targets of the RosettaAntibody benchmark (Figure S2). Although some neighboring residues were consistently attended to, we observed noticeable changes in attention patterns across the targets (Figure S3), demonstrating the sensitivity of the attention mechanism for identifying key interactions for a broad range of structures.

#### Repertoire sequence model learns evolutionary and structural representations

To better understand what properties of antibodies are accessible through unsupervised learning, we interrogated the representation learned by the LSTM encoder, which was trained only on sequences. First, we passed the entire set of paired heavy and light chain sequences from the OAS database through the network to generate embeddings like those used for the structure prediction model. The variable-length embedding for each sequence was averaged over its length to generate a fixed-size vector describing the entire sequence. We projected the vector embedding for each sequence into two dimensions via t-distributed stochastic neighbor embedding (t-SNE)^41^ and found that the sequences were naturally clustered by species (Figure 3D). Because the structural dataset is predominately composed of human and murine antibodies, the unsupervised features are likely providing evolutionary context that is otherwise unavailable.

The five non-H3 CDR loops typically adopt one of several canonical conformations^42,43^. Previous studies have identified distinct structural clusters for these loops and described each cluster by a characteristic sequence signature^44^. We hypothesized that our unsupervised learning model should detect these sequence signatures and thus encode information about the corresponding structural clusters. Similar to before, we created fixed-size embedding vectors for the five non-H3 loops by averaging the whole-sequence embedding over the residues of each loop (according to Chothia definitions^42^). In Figure 3E, we show t-SNE embeddings for the CDR1 loops labeled by their structural clusters from PyIgClassify^44^. These loops are highlighted because they have the most uniform class balance among structural clusters; similar plots for the remaining loops are provided in Figure S4. We observed clustering of labels for both CDR1 loops, indicating that the unsupervised model has captured some structural features of antibodies through sequence alone.

### Applicability to antibody design

Moving towards the goal of antibody design, we sought to test our method’s ability to distinguish between beneficial and disruptive mutations. First, we gathered a previously published deep mutational scanning (DMS) dataset for an anti-lysozyme antibody^45^. Anti-lysozyme was an ideal subject for evaluating our network’s design capabilities, as it was part of the benchmark set and thus already excluded from training. In the DMS dataset, anti-lysozyme was subjected to mutational scanning at 135 positions across the F_V_, including the CDR loops and the heavy-light chain interface. Each variant was transformed into yeast and measured for binding enrichment over the wild type.

#### Prediction confidence is indicative of mutation tolerability

We explored two strategies for evaluating mutations with our network. First, we measured the change in the network’s structure prediction confidence for a variant sequence relative to the wild type (visualized in Figure 4A) as a change in categorical cross entropy:

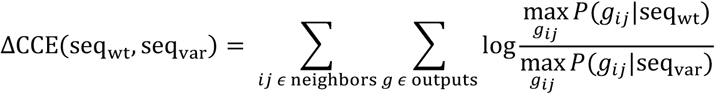

where seq_wt_ and seq_var_ are the wild type and variant sequences, respectively, and the conditional probability term describes the probability of a particular geometric output 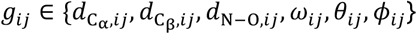 given seq_wt_ or seq_var_. Only residue pairs *ij* with predicted *d*_*Ca*_ < 10 Å were used in the calculation. Second, we used the LSTM decoder described previously to calculate the negative log likelihood of a particular point mutation given the wild type sequence, termed dLSTM:

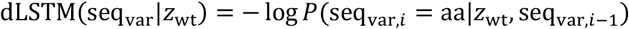

where seq_var_ is a variant sequence with a point mutation to *aa* at position *i*, and *z*_wt_ is the bi-LSTM encoder summary vector for the wild type sequence. To evaluate the discriminative power of the two metrics, we calculated ΔCCE and dLSTM for each variant in the anti-lysozyme dataset. We additionally calculated a combined metric as ΔCCE + 0.01 × dLSTM, roughly equating the magnitudes of both terms, and compared to the experimental binding data (Figure 4B). Despite having no explicit knowledge of the antigen, the network was moderately predictive of experimental binding enrichment (Figure 4C). The most successful predictions (true positives in Figure 4B) were primarily for mutations in CDR loop residues (Figure 4D). This is not surprising, given that our network has observed the most diversity in these hypervariable regions and is likely less calibrated to variance among framework residues. Nevertheless, if the ΔCCE + 0.01 × dLSTM were for ranking, all the top-8 and 22 of the top-100 single-point mutants identified would have experimental binding enrichments above the wild type (Figure 4E).

**Figure 4.**
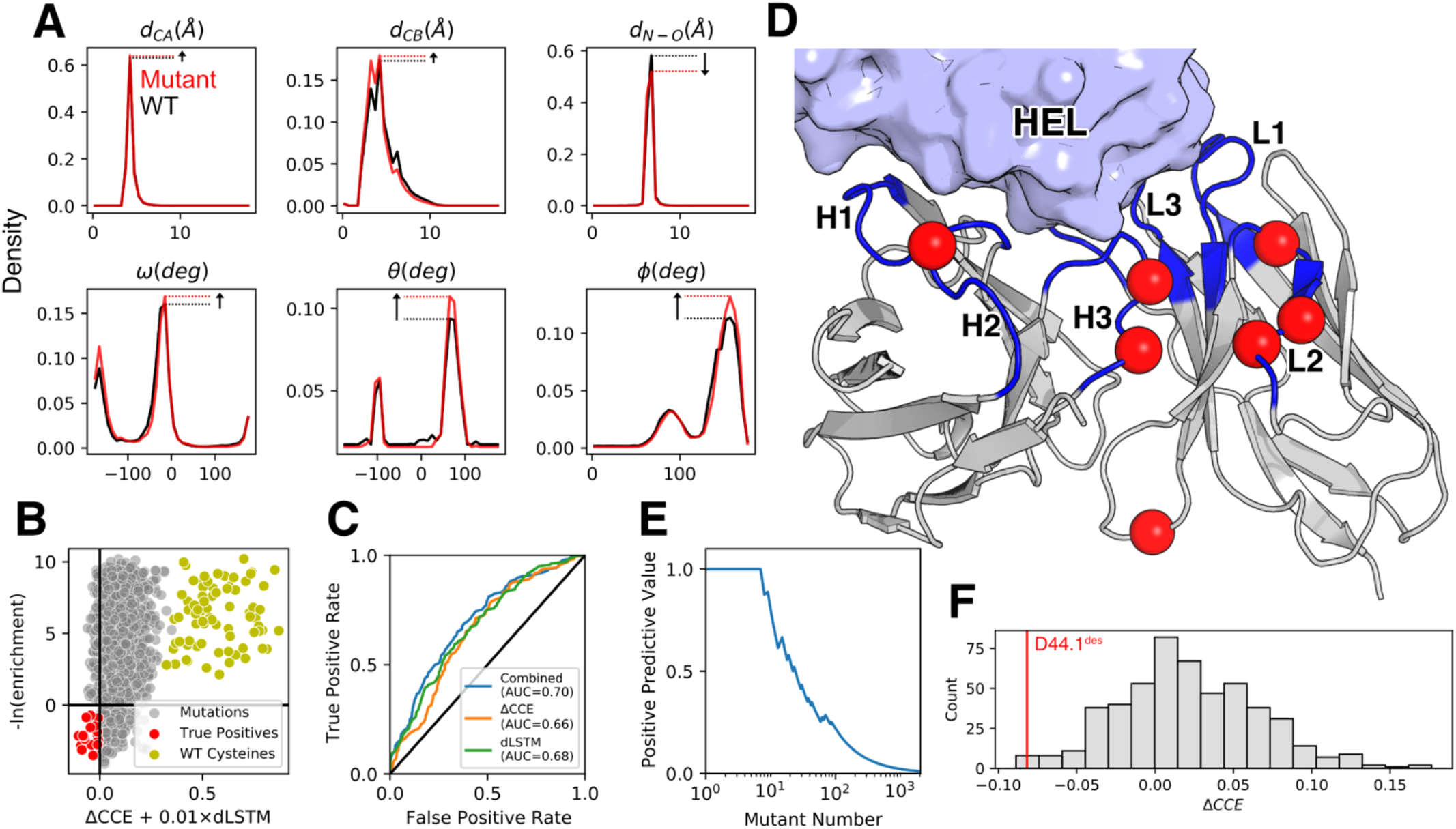
Prediction of mutational effects with DeepAb model. (A) Diagram of ΔCCE calculation for model output predictions for an arbitrary residue pair. Plots show the change in probability density of the predicted geometries for the residue pair after making a mutation. (B) Plot of the combined network metric against experimental binding enrichment over wild type, with negative values corresponding to beneficial mutations for both axes. True positive predictions (red) and mutations to wild type cysteines (yellow) are highlighted. (C) Receiver operating characteristic for predicting experimental binding enrichment over wild type with the combined network metric and each component metric. Area under the curve (AUC) values are provided for each metric. (D) Position of true positive predictions on anti-lysozyme F_V_ structure. (E) Positive predictive value for mutants ranked by the combined metric. (F) Comparison of ΔCCE for a designed eight-point variant (D44.1^des^, red) to sequences with random mutations at the same positions.

#### Network distinguishes stability-enhanced designs

The anti-lysozyme DMS dataset was originally assembled to identify residues for design of multi-point variants^45^. The authors designed an anti-lysozyme variant with eight mutations, called D44.1^des^, that displayed improved thermal stability and nearly tenfold increase in affinity. To determine whether our network could recognize the cumulative benefits of multiple mutations, we created a set of variants with random mutations at the same positions. We calculated ΔCCE for D44.1^des^ and the random variants and found that the model successfully distinguished the design (Figure 4F). We found similar success at distinguishing additional designs from the same publication (Figure S5). Despite being trained only for structure prediction, these results suggest that our model may be a useful tool for screening or ranking candidates in antibody design pipelines.

## Discussion

The results presented in this work build on advances in general protein structure prediction to effectively predict antibody F_V_ structures. We found that our deep learning method consistently produced more accurate structures than grafting-based alternatives on benchmarks of challenging, therapeutically relevant targets. As deep learning methods continue to improve, model interpretability will become increasingly important to ensure practitioners can gain insights beyond the primary predictive results. In addition to producing accurate structures, our method also provides interpretable insights into its predictions. Through the attention mechanism, we can track the network’s focus while predicting F_V_ structures. We demonstrated interpretation of predictions for a CDR H3 loop and identified several interactions with neighboring residues that the model deemed important for structure. In the future, similar insights could be used to guide antibody engineering efforts.

As part of this work, we developed an unsupervised representation model for antibody sequences. We found that critical features of antibody structure, including non-H3 loop clusters, were accessible through a simple LSTM encoder-decoder model. While we limited training to known pairs of heavy and light chains, several orders of magnitude more unpaired immunoglobins have been identified through next-generation repertoire sequencing experiments^28^. We anticipate that a more advanced language model trained on this larger sequence space will enable further advances across all areas of antibody bioinformatics research.

Deep learning models for antibody structure prediction present several promising avenues towards antibody design. In this work, we demonstrated how our network could be used to suggest or screen point mutations. Even with no explicit knowledge of the antigen, this approach was already moderately predictive of mutational tolerability. Inclusion of antigen structural context through extended deep learning models or traditional approaches like Rosetta should only improve these results. Other quantities of interest such as stability or developability metrics could be predicted by using the DeepAb network for transfer learning or feature engineering^12^. Furthermore, comparable networks for general protein structure prediction have recently been re-purposed for design through direct sequence optimization^46–48^. With minimal modification, our network should enable similar methods for antibody design.

## Methods

### Independent test sets

To evaluate the performance of our method, we considered two independent test sets. The first is the RosettaAntibody benchmark set of 49 structures, which was previously assembled to evaluate methods over a broad range of CDR H3 loop lengths (ranging 7-17 residues)^8,35^. Each structure in this set has greater than 2.5 Å resolution, a maximum R value of 0.2, and a maximum B factor of 80 Å². The second comes from a set of 56 clinical-stage antibody therapeutics with solved crystal structures, which was previously assembled to study antibody developability^36^. We removed five of the therapeutic antibodies that were missing one or more CDR loops (PDB IDs: 3B2U, 3C08, 3HMW, 3S34, and 4EDW) to create a therapeutic benchmark set. The two sets shared two common antibodies (PDB IDs: 3EO9 and 3GIZ) that we removed from the therapeutic benchmark set.

While benchmarking alternative methods, we found that some methods were unable to produce structures for every target. To compare consistently across all methods, we report values for only the targets that all methods succeeded in modeling. However, we note that DeepAb was capable of producing structures for all of the targets attempted. From the RosettaAntibody benchmark set we omit PDB IDs 1X9Q and 3IFL. From the therapeutic benchmark set we omit PDB IDs 4D9Q, 4K3J, 4O02, and 5VKK. We additionally omit the long L3 loop of target 3MLR, which not all alternative methods were able to model. In total, metrics are reported for 92 targets: 47 from the RosettaAntibody benchmark and 45 from the therapeutic benchmark. We use the Chothia CDR loop definitions to measure RMSD throughout this work^42^.

### Representation learning on repertoire sequences

#### Training dataset

To train the sequence model, paired F_V_ heavy and light chain sequences were collected from the Observed Antibody Space (OAS) database^28^, a set of immunoglobin sequences from next-generation sequencing experiments of immune repertoires. Each sequence in the database had previously been parsed with ANARCI^49^ to annotate sequences and detect potentially erroneous entries. Sequences indicated to have failed ANARCI parsing were discarded from the training dataset. We additionally remove any redundant sequences. These steps resulted in a set of 118,386 sequences for model training.

#### Model and training details

To learn representations of immunoglobin sequences, we adopted an RNN encoder-decoder model^26^ consisting of two LSTMs^27^. In an encoder-decoder model, the encoder learns to summarize the input sequence into a fixed-dimension summary vector, from which the decoder learns to reconstruct the original sequence. For the encoder model, we used a bidirectional two-layer stacked LSTM with a hidden state size of 64. The model input was created by concatenation of paired heavy and light chain sequences to form a single sequence. Three additional tokens were added to the sequence to mark the beginning of the heavy chain, the end of the heavy chain, and the end of the light chain. The concatenated sequence was one-hot encoded, resulting in an input of dimension (*L* + 3) × 23, where *L* is the combined heavy and light chain length. The summary vector is generated by stacking the final hidden states from the forward and backward encoder LSTMs, followed by a linear transformation from 128 to 64 dimensions and *tanh* activation. For the decoder model, we used a two-layer stacked LSTM with a hidden state size of 64. The decoder takes as input the summary vector and the previously decoded amino acid to sequentially predict the original amino acid sequence.

The model was trained using cross-entropy loss and the Adam optimizer^50^ with a learning rate of 0.01. A teacher forcing rate of 0.5 was used to stabilize training. The model was trained on one NVIDIA K80 GPU, requiring ∼4 hours for 5 epochs over the entire dataset. We used a batch size of 128, maximized to fit into GPU memory.

### Predicting inter-residue geometries from antibody sequence

#### Training dataset

To train the structure prediction model, we collected a set of F_V_ structures from the Structural Antibody Database (SAbDab)^32^, a curated set of antibody structures from the PDB^51^. We removed structures with less than 4 Å resolution and applied a 99% sequence identity threshold to remove redundant sequences. We chose this high sequence similarity due to the high conservation characteristic of antibody sequences. Finally, any targets from the benchmark sets, or structures with 99% sequence similarity to a target, were removed from the training dataset. These steps resulted in a set of 1,692 F_v_ structures for model training.

#### Model and training details

The structure prediction model takes as input concatenated heavy and light chain sequences. The sequences are one-hot encoded and passed through two parallel branches: a 1D ResNet and the bi-LSTM encoder described above. For the 1D ResNet, we add an additional delimiter channel to mark the end of the heavy chain, resulting in a dimension of *L* × 21, where *L* is the combined heavy and light chain length. The 1D ResNet begins with a 1D convolution that projects the input features up to dimension *L* × 64, followed by three 1D ResNet blocks (two 1D convolutions with kernel size 17) that maintain dimensionality. The second branch consists of the pre-trained bi-LSTM encoder. Before passing the one-hot encoded sequence to the bi-LSTM, we add the three delimiters described previously, resulting in dimension (*L* + 3) × 23. From the bi-LSTM, we concatenate the hidden states from the forward and backward LSTMs after encoding each residue, resulting in dimension *L* × 128. The outputs of the 1D ResNet and the bi-LSTM are stacked to form a final sequential tensor of dimension *L* × 160. We transform the sequential tensor to pairwise data by concatenating row-and column-wise expansions. The pairwise data, dimension *L* × *L* × 320, is passed to the 2D ResNet. The 2D ResNet begins with a 2D convolution that reduces dimensionality to *L* × *L* × 64, followed by 25 2D ResNet blocks (two 2D convolutions with kernel size 5 × 5) that maintain dimensionality. The 2D ResNet blocks cycle through convolution dilation values of 1, 2, 4, 8, and 16 (five cycles in total). After the 2D ResNet, the network branches into six separate paths. Each output branch consists of a 2D convolution that projects down to dimension *L* × *L* × 37, followed by a recurrent criss-cross attention (RCCA) module^39^. The RCCA modules use two criss-cross attention operations that share weights, allowing each residue pair to gather information across the entire spatial dimension. Attention queries and keys are projected to dimension *L* × *L* × 1 (one attention head). Symmetry is enforced for 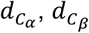, and *ω* predictions by averaging the final outputs with their transposes. All convolutions in the network are followed by ReLU activation. In total, the model contains about 6.4 million trainable parameters.

We trained five models on random 90/10% training/validation splits and averaged over model logits to make predictions, following previous methods^8,19^. Models were trained using focal loss^30^ and the Adam optimizer^50^ with a learning rate of 0.01. Learning rate was reduced upon plateauing of the validation loss. Each model was trained on one NVIDIA K80 GPU, requiring ∼60 hours for 60 epochs over the entire dataset.

### Structure realization

#### Multi-dimensional scaling

From the network predictions, we create real-value matrices for the 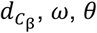, and *ϕ* outputs by taking the midpoint value of the modal probability bin for each residue pair. From these real-valued distances and orientations, we create an initial backbone atom (N, C_α_, and C) distance matrix. For residue pairs predicted to have 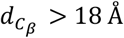, we approximate the distances between atoms using the Floyd-Warshall shortest path algorithm^52^. From this distance matrix, we use multi-dimensional scaling (MDS)^53^ to produce an initial set of 3D coordinates. The initial structures from MDS typically contained atom clashes and non-ideal geometries that required further refinement.

#### Energy minimization refinement

Initial structures from MDS were refined by constrained energy minimization in Rosetta. For each pair of residues, the predicted distributions for each output were converted to energy potentials by negating the raw model logits (*i*.*e*. without softmax activation) and dividing by the squared 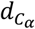 prediction. The discrete potentials were converted to continuous functions using the built-in Rosetta spline function. We disregarded potentials for residue pairs with predicted 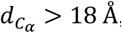, as well as those with a modal bin probability below 10%. For *d*_*N*−*O*_ potentials, we also discarded with predicted *d*_*N*−*O*_ > 5 Å or modal bin probability below 30% to create a local backbone hydrogen-bonding potential. The remaining potentials are applied to the MDS structure as inter-residue constraints in Rosetta.

Modeling in Rosetta begins with a coarse-grained representation, in which the side-chain atoms are represented as a single artificial atom (centroid). The centroid model is optimized by gradient-based energy minimization (*MinMover*) using the L-BFGS algorithm^33,34^. After centroid optimization, we add side-chain atoms and relax the structure to reduce steric clashes (*FastRelax*). Finally, we repeat the gradient-based energy minimization step in the full-atom representation to produce a final model. We repeat this procedure to produce 50 decoy models and select the structure with the lowest energy as the final prediction. Only the relaxation step in the protocol is non-deterministic, leading to high convergence among decoys. In practice, we expect 5-10 decoys will be sufficient for most applications.

### Predicting structures with other recent methods

To contextualize the performance of our method, we benchmarked three recent methods for antibody F_V_ structure prediction: RosettaAntibody-G^6^, RepertoireBuilder^5^, and ABodyBuilder^3^. RosettaAntibody-G predictions were generated using the command-line arguments recommended by Jeliazkov et al^6^ (Appendix S1). We note that we only used the RosettaAntibody grafting protocol (*antibody*), omitting the extensive but time-consuming H3 loop sampling (*antibody_H3*)^4,6^. RepertoireBuilder and ABodyBuilder predictions were generated using their respective web servers. For each target in the benchmarks, we excluded structures with sequence similarity greater than 99% from use for predictions, to mirror the conditions of our training set. We note that this sequence cutoff does not prevent methods from grafting identical loops from slightly different sequences.

### Attention matrix calculation

During the criss-cross attention operation^39^, we create an attention matrix ***A****∈*ℝ^*L*×*L*×(2*L*−1)^, where for each residue pair in the *L* × *L* spatial dimension we have 2*L* − 1 entries corresponding to the attention values over other residue pairs in the same row and column (including the residue pair itself). To interpret the total attention between pairs of residues, we simplify the attention matrix to ***A***^′^*∈*ℝ^*L*×(2*L*−1)^, where for each residue *i* in the sequence we only consider the attention values in the *i*-th row and column. In ***A***′, for each residue *i* there are two attention values for each other residue *j*, corresponding to the row- and column-wise attention between *i* and *j*. We further simplify by summing these row- and column-wise attention values, resulting in an attention matrix ***A***^′′^*∈*ℝ^*L*×*L*^, containing the total attention between pairs of residues. In the main text, we refer to ***A***′′ as ***A*** for simplicity.

## Acknowledgements

We thank Dr. Sai Pooja Mahajan for helpful discussions and advice. This work was supported by National Institutes of Health grants R01-GM078221 and T32-GM008403 (J.A.R.) and AstraZeneca (J.A.R.). Computational resources were provided by the Maryland Advanced Research Computing Cluster (MARCC).

## Data Availability

The source code for inter-residue geometry prediction and structure realization with Rosetta, as well as pretrained models, are available at github.com/rosettacommons/deepab. The structures predicted by DeepAb and alternative methods for benchmarking will be made available prior to publication.

