## Supplementary Material for "Antibody structure prediction using interpretable deep learning"

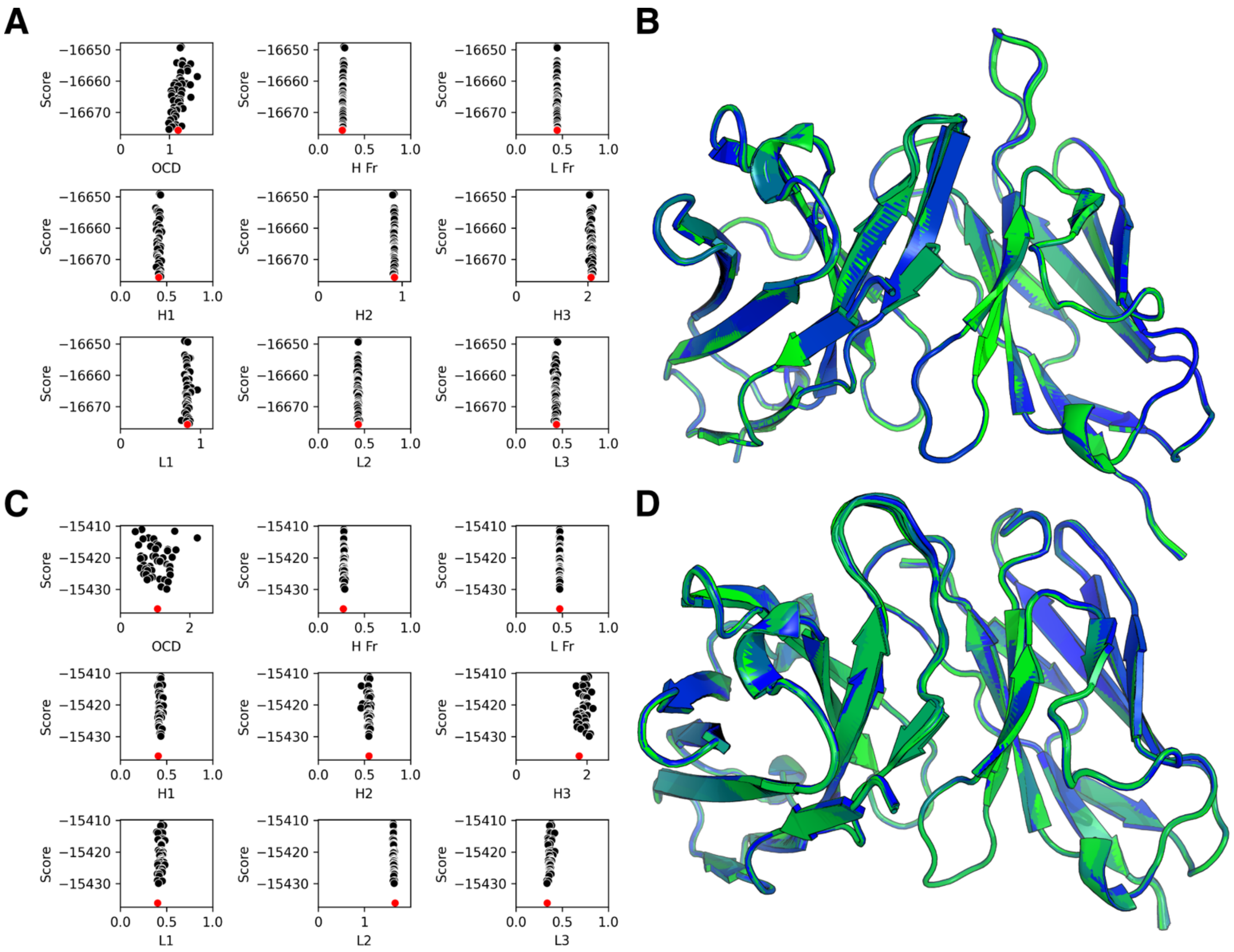


**Figure S1. Convergence of predicted structures for two benchmark examples.** (A) Funnel plots showing accuracy (OCD, RMSD) versus score for 50 DeepAb decoys for target 3PP3 (therapeutic benchmark), with low-scoring structure in red. (B) Superimposed decoy structures for target 3PP3. (C) Funnel plots showing accuracy (OCD, RMSD) versus score for 50 DeepAb decoys for target 3I9G (RosettaAntibody benchmark), with low-scoring structure in red. (D) Superimposed decoy structures for target 3I9G.

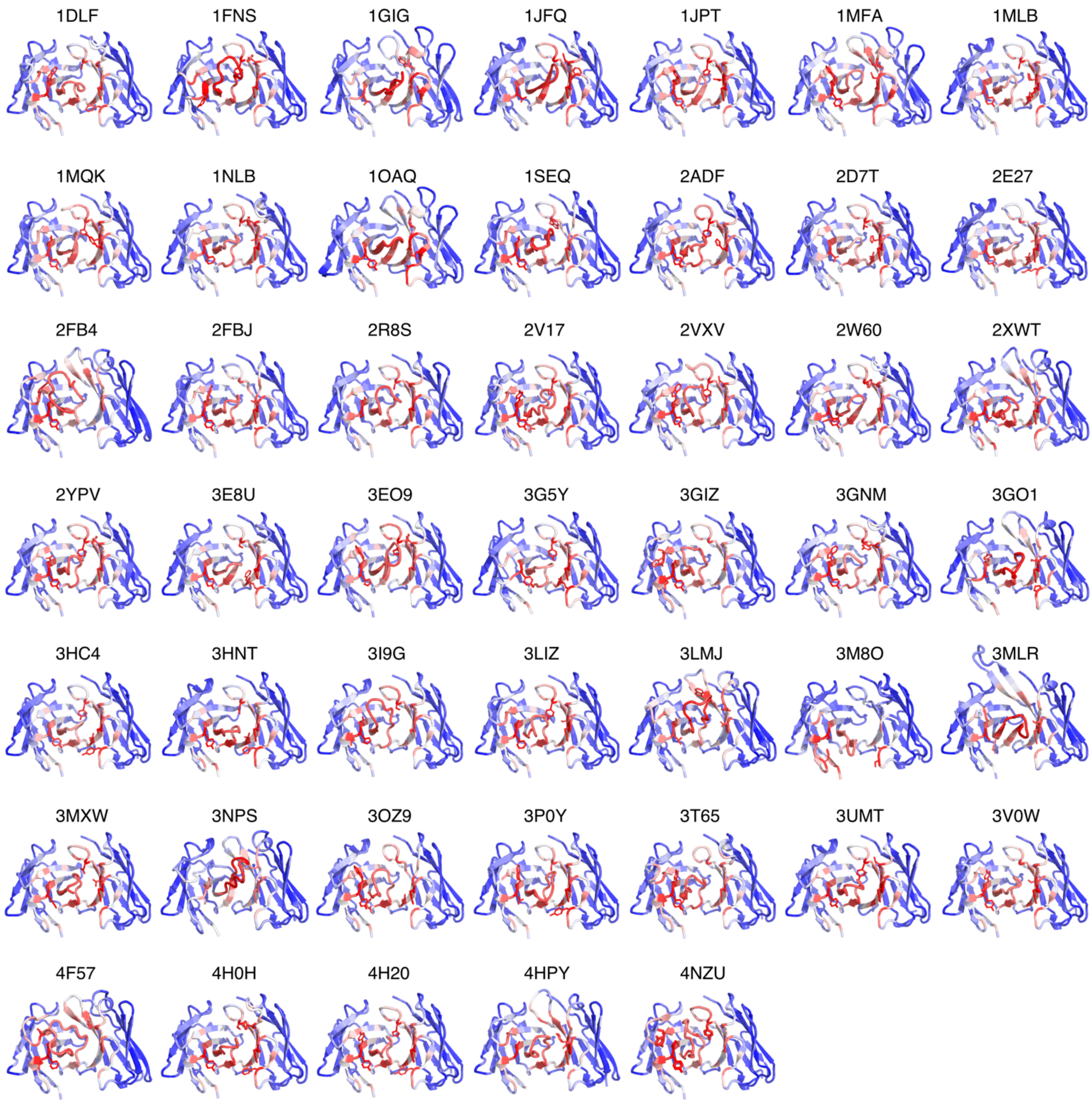


**Figure S2. H3 loop attention for RosettaAntibody benchmark targets.** Model $C_\alpha$ attention while predicting H3 loop structures for each of the 47 targets in the RosettaAntibody benchmark. Attention values increase from blue to red. For each target, the side chains of the five most attended non-H3 residues are represented as sticks.

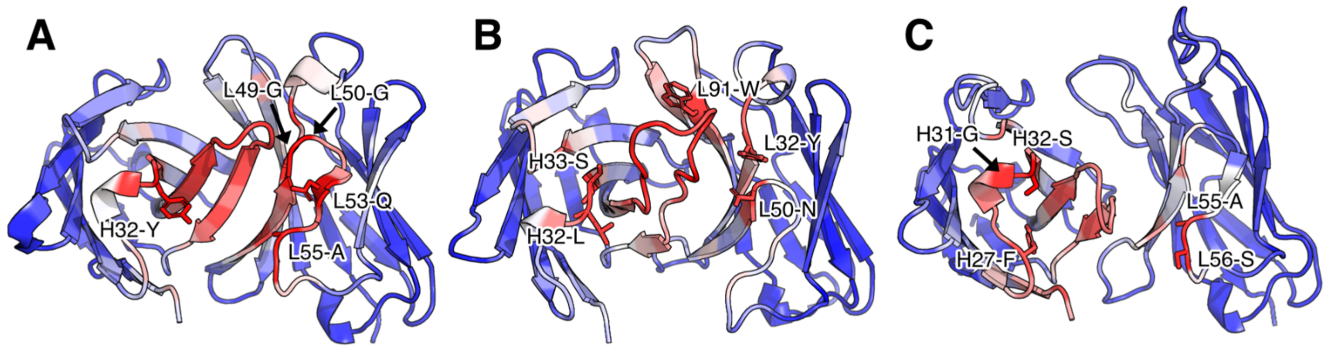


**Figure S3. Variability of key residues identified by attention mechanism.** Model $C_\alpha$ attention while predicting H3 loop structures for three targets in the RosettaAntibody benchmark. Attention values increase from blue to red. For each target, the side chains of the five most attended non-H3 residues are represented as sticks. (A) H3 attention for 1OAQ prediction. (B) H3 attention for 3MLJ prediction. (C) H3 attention for 3M8O prediction.

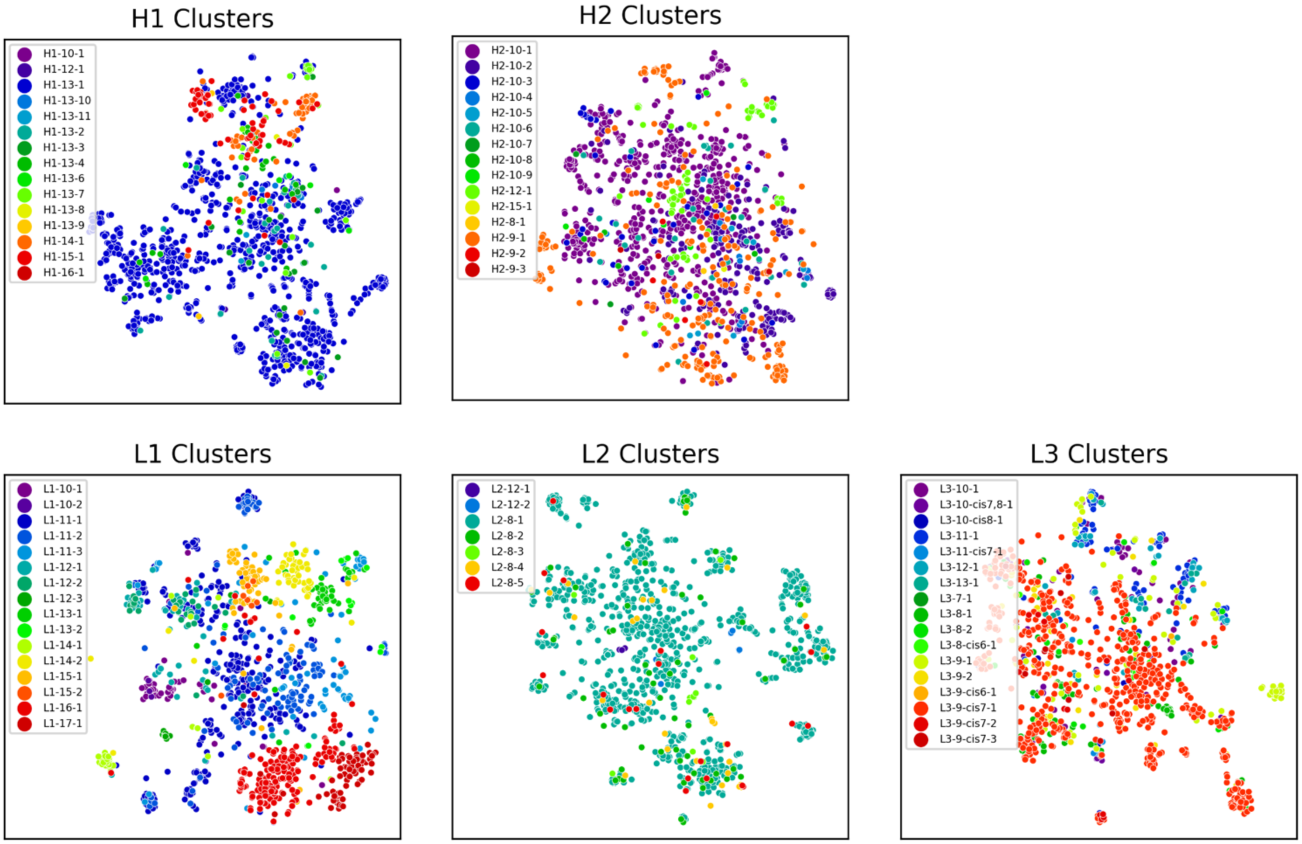


**Figure S4. Non-H3 CDR loop t-SNE embeddings labeled by structural clusters.** CDR-specific embeddings are created by averaging the bi-LSTM encoder hidden states of residues for each CDR loop.

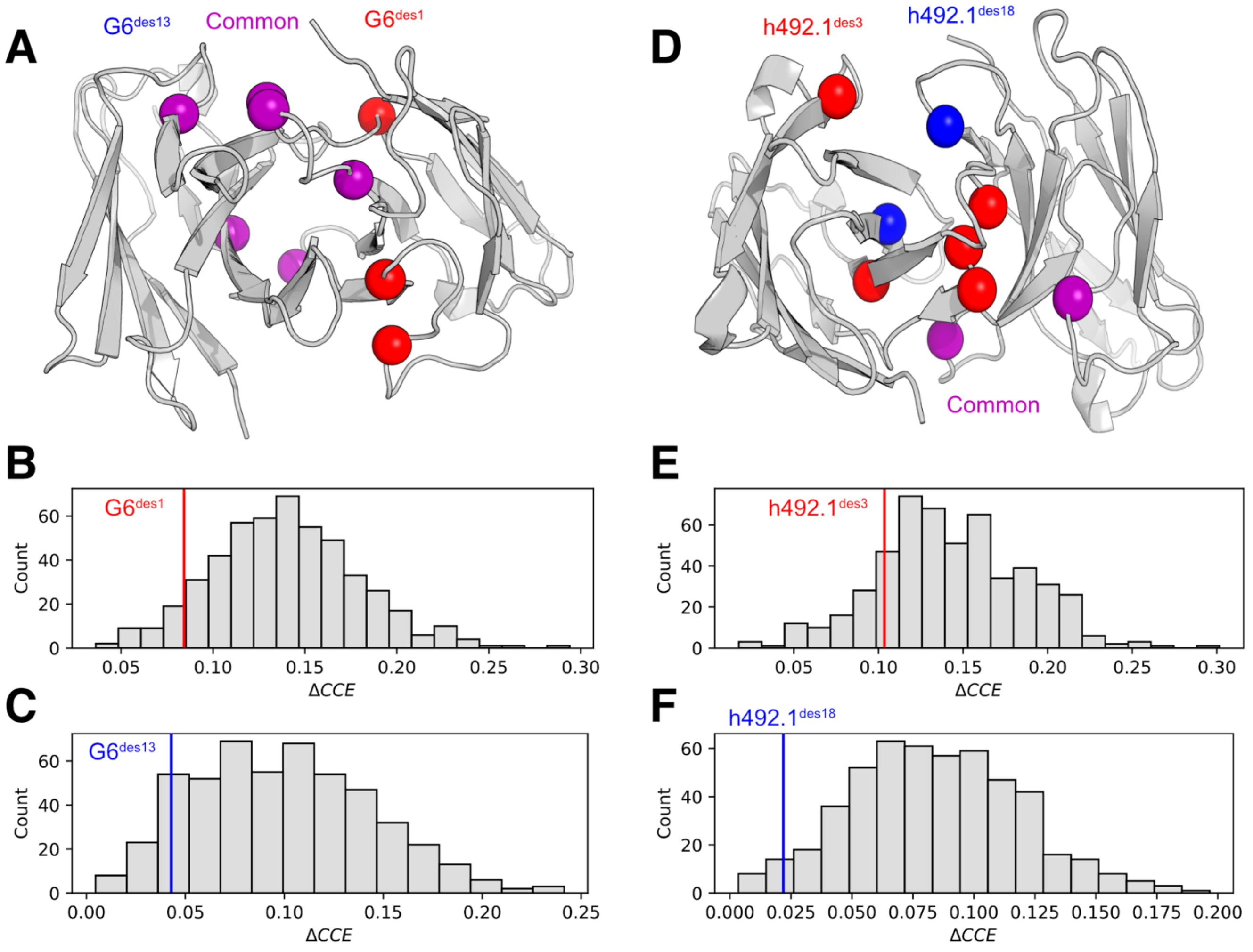


**Figure S5. Identification of stable multi-point variants for two AbLIFT designs.** Both wild type structures were present in the training dataset, resulting in a slight bias for the native sequence. (A) Mutation positions for two anti-VEGF multi-point variants presented by Warszawski *et al*[2]. (B) Comparison of $\Delta CCE$ values for G6[des1] (nine-point variant) and random nine-point variants at the same positions. (C) Comparison of $\Delta CCE$ values for G6[des13] (six-point variant) and random six-point variants at the same positions. (D) Mutation positions for two anti-QSOX1 multi-point variants. (E) Comparison of $\Delta CCE$ values for h492.1[des3] (seven-point variant) and random seven-point variants at the same positions. (F) Comparison of $\Delta CCE$ values for h492.1[des18] (four-point variant) and random four-point variants at the same positions.

**Table S1. CDR loop RMSD and OCD results for DeepAb on RosettaAntibody benchmark.**

| Target | OCD | H Fr (Å) | H1 (Å) | H2 (Å) | H3 (Å) | L Fr (Å) | L1 (Å) | L2 (Å) | L3 (Å) |
|---|---|---|---|---|---|---|---|---|---|
| 1dlf | 6.04 | 0.65 | 0.92 | 0.82 | 3.84 | 0.51 | 0.37 | 0.64 | 0.32 |
| 1fns | 1.72 | 0.37 | 0.62 | 1.15 | 2.01 | 0.32 | 0.51 | 0.27 | 0.51 |
| 1gig | 1.95 | 0.39 | 0.64 | 0.47 | 2.49 | 0.47 | 0.36 | 0.99 | 1.14 |
| 1jfq | 3.20 | 0.49 | 0.49 | 1.07 | 1.10 | 0.33 | 0.43 | 0.21 | 0.54 |
| 1jpt | 1.42 | 0.43 | 0.36 | 0.67 | 1.29 | 0.24 | 0.72 | 0.41 | 0.37 |
| 1mfa | 2.08 | 0.45 | 0.50 | 0.69 | 1.45 | 0.29 | 0.89 | 0.41 | 1.24 |
| 1mlb | 5.36 | 0.46 | 0.42 | 0.60 | 1.14 | 0.42 | 0.44 | 0.76 | 0.64 |
| 1mqk | 2.68 | 0.24 | 0.29 | 0.34 | 1.04 | 0.48 | 0.46 | 0.38 | 1.76 |
| 1nlb | 3.21 | 0.27 | 0.21 | 0.59 | 0.56 | 0.50 | 0.77 | 0.33 | 0.35 |
| 1oaq | 1.33 | 0.39 | 0.37 | 0.67 | 1.93 | 0.38 | 0.58 | 0.39 | 0.40 |
| 1seq | 2.45 | 0.39 | 0.66 | 0.58 | 2.93 | 0.36 | 0.41 | 0.20 | 0.40 |
| 2adf | 2.87 | 0.42 | 0.48 | 0.61 | 1.77 | 0.53 | 0.75 | 0.28 | 0.59 |
| 2d7t | 15.91 | 0.65 | 0.80 | 0.78 | 2.27 | 0.44 | 0.42 | 0.36 | 1.00 |
| 2e27 | 3.29 | 0.34 | 0.54 | 0.53 | 4.30 | 0.58 | 0.71 | 0.49 | 1.90 |
| 2fb4 | 5.83 | 0.34 | 0.30 | 0.34 | 3.53 | 0.88 | 0.91 | 0.47 | 0.63 |
| 2fbj | 6.73 | 0.46 | 0.57 | 1.01 | 1.44 | 0.45 | 1.02 | 0.62 | 1.99 |
| 2r8s | 2.59 | 0.48 | 0.64 | 1.83 | 2.55 | 0.37 | 0.32 | 0.41 | 0.49 |
| 2v17 | 2.12 | 0.36 | 0.44 | 1.26 | 2.44 | 0.67 | 0.69 | 0.35 | 0.86 |
| 2vxv | 1.95 | 0.35 | 0.65 | 1.09 | 5.28 | 0.35 | 0.26 | 0.90 | 1.77 |
| 2w60 | 4.64 | 0.29 | 0.36 | 0.54 | 1.97 | 0.36 | 0.43 | 0.30 | 0.49 |
| 2xwt | 0.46 | 0.43 | 1.24 | 0.54 | 3.13 | 0.32 | 0.61 | 1.07 | 0.99 |
| 2ypv | 3.73 | 0.82 | 0.69 | 0.70 | 2.36 | 0.33 | 0.68 | 0.39 | 0.55 |
| 3e8u | 3.26 | 0.48 | 0.56 | 0.59 | 1.30 | 0.37 | 0.51 | 0.42 | 0.35 |
| 3eo9 | 4.42 | 0.36 | 0.53 | 0.68 | 2.42 | 0.34 | 0.41 | 0.39 | 0.79 |
| 3g5y | 3.75 | 0.29 | 0.52 | 0.37 | 0.72 | 0.60 | 0.24 | 0.35 | 0.42 |
| 3giz | 2.10 | 0.37 | 0.56 | 0.43 | 1.95 | 0.31 | 0.25 | 0.28 | 0.25 |
| 3gnm | 0.81 | 0.29 | 0.44 | 0.30 | 2.78 | 0.40 | 0.63 | 0.19 | 0.47 |
| 3go1 | 2.13 | 0.36 | 1.47 | 1.22 | 3.28 | 0.47 | 0.95 | 0.83 | 1.83 |
| 3hc4 | 4.13 | 1.02 | 0.49 | 0.99 | 0.89 | 0.41 | 0.46 | 0.30 | 0.37 |
| 3hnt | 3.28 | 0.39 | 0.19 | 0.66 | 1.19 | 0.49 | 0.36 | 0.90 | 0.93 |
| 3i9g | 1.24 | 0.27 | 0.41 | 0.55 | 1.78 | 0.47 | 0.40 | 1.66 | 0.34 |
| 3liz | 1.67 | 0.33 | 0.52 | 0.37 | 4.77 | 0.50 | 0.66 | 0.32 | 0.30 |
| 3lmj | 3.20 | 0.40 | 3.16 | 5.81 | 5.26 | 0.27 | 0.46 | 0.35 | 1.68 |
| 3m8o | 4.42 | 0.56 | 1.48 | 1.19 | 2.98 | 0.48 | 1.49 | 0.32 | 1.08 |
| 3mlr | 2.36 | 0.51 | 0.57 | 0.53 | 3.65 | 0.43 | 0.50 | 0.34 | - |
| 3mxw | 1.03 | 0.41 | 0.59 | 0.56 | 1.08 | 0.32 | 0.30 | 0.25 | 0.68 |
| 3nps | 21.84 | 1.19 | 3.24 | 1.76 | 4.84 | 0.45 | 0.67 | 0.48 | 1.01 |
| 3oz9 | 2.89 | 0.48 | 0.53 | 0.90 | 1.21 | 0.46 | 0.85 | 0.21 | 1.52 |
| 3p0y | 2.10 | 0.37 | 0.70 | 0.98 | 1.53 | 0.21 | 0.39 | 0.30 | 1.53 |
| 3t65 | 1.91 | 0.28 | 0.36 | 0.40 | 1.48 | 0.39 | 0.49 | 0.36 | 0.39 |
| 3umt | 2.52 | 0.36 | 0.60 | 0.73 | 2.05 | 0.47 | 0.79 | 0.47 | 0.30 |
| 3v0w | 4.87 | 0.42 | 0.86 | 0.53 | 0.91 | 0.44 | 0.34 | 0.33 | 0.44 |
| 4f57 | 2.22 | 0.27 | 0.32 | 1.08 | 1.36 | 0.23 | 0.34 | 0.23 | 0.73 |
| 4h0h | 3.06 | 0.30 | 0.33 | 0.62 | 1.22 | 0.47 | 0.35 | 0.37 | 0.29 |
| 4h20 | 3.81 | 0.52 | 0.26 | 0.60 | 2.63 | 0.35 | 0.34 | 0.29 | 0.83 |
| 4hpy | 3.71 | 0.33 | 2.58 | 0.29 | 1.94 | 0.27 | 0.66 | 0.34 | 2.23 |
| 4nzu | 4.35 | 0.31 | 0.27 | 0.73 | 5.64 | 0.40 | 0.26 | 0.42 | 0.37 |
| **Mean** | 3.67 | 0.43 | 0.72 | 0.85 | 2.33 | 0.42 | 0.55 | 0.45 | 0.86 |
| **SD** | 3.58 | 0.18 | 0.66 | 0.81 | 1.32 | 0.12 | 0.25 | 0.28 | 0.58 |

**Table S2. CDR loop RMSD and OCD results for DeepAb on therapeutic benchmark.**

| Target | OCD | H Fr (Å) | H1 (Å) | H2 (Å) | H3 (Å) | L Fr (Å) | L1 (Å) | L2 (Å) | L3 (Å) |
|---|---|---|---|---|---|---|---|---|---|
| 1bey | 1.64 | 0.58 | 1.45 | 0.86 | 2.35 | 0.62 | 0.73 | 0.55 | 0.65 |
| 1cz8 | 1.92 | 0.30 | 0.64 | 0.76 | 1.65 | 0.24 | 0.30 | 0.28 | 0.38 |
| 1mim | 1.60 | 0.49 | 0.54 | 0.49 | 1.01 | 0.36 | 0.61 | 0.48 | 0.81 |
| 1sy6 | 1.78 | 0.32 | 0.46 | 0.62 | 2.16 | 0.40 | 0.32 | 0.31 | 0.90 |
| 1yy8 | 1.78 | 0.98 | 0.49 | 0.68 | 0.62 | 0.42 | 0.57 | 0.49 | 0.74 |
| 2hwz | 3.17 | 0.25 | 0.50 | 0.35 | 1.93 | 0.46 | 0.70 | 0.37 | 2.40 |
| 3eo0 | 2.50 | 0.31 | 3.25 | 1.05 | 4.49 | 0.27 | 1.27 | 0.23 | 0.44 |
| 3gkw | 9.94 | 1.24 | 2.79 | 1.08 | 7.48 | 0.42 | 0.72 | 0.35 | 0.91 |
| 3nfs | 3.88 | 0.46 | 0.44 | 0.54 | 1.10 | 0.39 | 0.65 | 0.36 | 0.84 |
| 3o2d | 5.31 | 0.38 | 0.39 | 0.71 | 2.26 | 0.24 | 0.54 | 0.27 | 0.40 |
| 3pp3 | 1.18 | 0.26 | 0.42 | 0.91 | 2.09 | 0.45 | 0.84 | 0.43 | 0.44 |
| 3qwo | 2.37 | 0.32 | 0.65 | 1.24 | 1.58 | 0.27 | 0.39 | 0.43 | 0.54 |
| 3u0t | 9.53 | 0.58 | 1.60 | 0.79 | 3.07 | 0.50 | 0.83 | 0.54 | 1.01 |
| 4cni | 3.76 | 0.32 | 0.31 | 1.02 | 2.79 | 0.44 | 0.33 | 0.39 | 0.63 |
| 4dn3 | 2.61 | 0.40 | 2.29 | 0.79 | 1.82 | 0.35 | 1.27 | 0.34 | 1.26 |
| 4g5z | 3.98 | 0.30 | 0.36 | 0.44 | 1.45 | 0.47 | 0.30 | 0.57 | 0.46 |
| 4g6k | 5.89 | 0.30 | 0.40 | 1.05 | 1.87 | 0.46 | 0.73 | 0.27 | 1.06 |
| 4hkz | 4.03 | 0.25 | 0.43 | 0.46 | 3.04 | 0.26 | 0.27 | 0.28 | 0.56 |
| 4i77 | 3.24 | 0.41 | 0.89 | 1.04 | 2.37 | 0.29 | 0.80 | 0.27 | 0.60 |
| 4irz | 1.77 | 0.38 | 0.43 | 0.52 | 3.05 | 0.41 | 1.00 | 0.41 | 1.38 |
| 4kaq | 3.63 | 0.54 | 0.45 | 0.58 | 2.38 | 0.37 | 0.49 | 0.32 | 0.38 |
| 4m6n | 6.05 | 0.35 | 0.70 | 0.40 | 2.76 | 0.28 | 1.32 | 0.29 | 2.02 |
| 4nyl | 5.31 | 0.52 | 0.35 | 0.56 | 4.74 | 0.55 | 0.52 | 0.46 | 0.78 |
| 4od2 | 1.38 | 0.44 | 0.42 | 0.61 | 5.53 | 0.45 | 0.78 | 1.36 | 2.14 |
| 4ojf | 1.35 | 0.36 | 0.28 | 1.09 | 1.71 | 0.24 | 0.42 | 0.32 | 0.55 |
| 4qxg | 0.95 | 0.37 | 0.37 | 0.47 | 1.55 | 0.36 | 0.44 | 0.46 | 1.67 |
| 4x7s | 1.99 | 0.37 | 1.43 | 0.65 | 2.18 | 0.29 | 0.39 | 0.22 | 0.55 |
| 4ypg | 3.94 | 0.38 | 0.62 | 0.45 | 3.00 | 0.37 | 0.45 | 0.66 | 0.86 |
| 5csz | 5.58 | 0.23 | 0.34 | 0.38 | 2.11 | 0.30 | 0.89 | 0.36 | 0.98 |
| 5dk3 | 4.00 | 0.32 | 0.54 | 0.58 | 2.49 | 0.33 | 0.32 | 0.55 | 0.52 |
| 5ggq | 2.13 | 0.31 | 0.34 | 0.54 | 1.45 | 0.28 | 0.18 | 0.31 | 0.20 |
| 5ggu | 5.51 | 0.24 | 0.43 | 0.46 | 6.14 | 0.32 | 0.27 | 0.25 | 0.34 |
| 5i5k | 8.73 | 0.73 | 0.63 | 0.79 | 4.57 | 0.84 | 0.78 | 0.64 | 0.95 |
| 5jxe | 3.98 | 0.29 | 0.67 | 0.46 | 2.85 | 0.59 | 0.49 | 0.58 | 0.77 |
| 5kmv | 6.17 | 0.52 | 0.62 | 0.71 | 0.73 | 0.34 | 0.58 | 0.45 | 0.51 |
| 5l6y | 3.89 | 0.25 | 0.47 | 0.54 | 5.44 | 0.21 | 0.28 | 0.52 | 2.29 |
| 5n2k | 2.78 | 0.24 | 0.35 | 0.29 | 2.70 | 0.35 | 0.51 | 0.32 | 4.55 |
| 5nhw | 2.43 | 0.29 | 0.69 | 0.48 | 0.80 | 0.26 | 0.76 | 0.37 | 4.34 |
| 5sx4 | 0.46 | 0.46 | 0.94 | 0.69 | 0.88 | 0.33 | 0.26 | 0.34 | 0.57 |
| 5tru | 3.00 | 0.33 | 0.40 | 0.43 | 1.34 | 0.30 | 1.01 | 0.25 | 1.22 |
| 5vh3 | 3.95 | 0.31 | 0.67 | 0.36 | 0.95 | 0.40 | 0.32 | 0.38 | 0.42 |
| 5wuv | 2.83 | 0.43 | 0.69 | 0.87 | 2.82 | 0.24 | 0.22 | 0.32 | 0.56 |
| 5xxy | 2.99 | 0.38 | 0.54 | 0.94 | 1.30 | 0.37 | 1.10 | 0.56 | 0.44 |
| 5y9k | 1.98 | 0.38 | 2.29 | 0.98 | 2.07 | 0.29 | 0.72 | 0.63 | 1.21 |
| 6and | 1.56 | 0.33 | 0.58 | 0.80 | 2.57 | 0.32 | 0.52 | 0.38 | 0.51 |
| **Mean** | 3.52 | 0.40 | 0.77 | 0.68 | 2.52 | 0.37 | 0.60 | 0.42 | 1.02 |
| **SD** | 2.16 | 0.19 | 0.67 | 0.24 | 1.50 | 0.12 | 0.30 | 0.19 | 0.92 |

**Table S3. CDR loop RMSD and OCD results for RosettaAntibody-G on RosettaAntibody benchmark.**

| Target | OCD | H Fr (Å) | H1 (Å) | H2 (Å) | H3 (Å) | L Fr (Å) | L1 (Å) | L2 (Å) | L3 (Å) |
|---|---|---|---|---|---|---|---|---|---|
| 1dlf | 9.61 | 0.69 | 1.17 | 1.14 | 4.92 | 0.61 | 0.67 | 1.00 | 0.64 |
| 1fns | 3.22 | 0.58 | 0.79 | 1.20 | 4.32 | 0.35 | 0.67 | 0.44 | 1.13 |
| 1gig | 8.70 | 0.59 | 0.68 | 0.86 | 4.35 | 0.90 | 0.78 | 2.29 | 1.00 |
| 1jfq | 5.07 | 0.67 | 0.97 | 1.07 | 1.34 | 0.51 | 0.40 | 0.83 | 0.74 |
| 1jpt | 6.69 | 0.52 | 1.62 | 0.99 | 1.48 | 0.30 | 0.67 | 0.90 | 0.75 |
| 1mfa | 7.30 | 0.76 | 0.82 | 1.07 | 3.39 | 0.93 | 2.37 | 0.98 | 1.29 |
| 1mlb | 4.15 | 0.59 | 1.23 | 0.58 | 5.45 | 0.59 | 0.54 | 1.18 | 0.85 |
| 1mqk | 6.21 | 0.41 | 0.71 | 0.88 | 1.78 | 0.71 | 0.79 | 0.81 | 1.95 |
| 1nlb | 4.41 | 0.52 | 0.62 | 0.52 | 1.47 | 0.55 | 0.54 | 0.79 | 0.89 |
| 1oaq | 7.46 | 0.76 | 1.22 | 0.85 | 1.96 | 0.88 | 0.64 | 1.72 | 1.21 |
| 1seq | 2.73 | 0.46 | 1.04 | 1.61 | 6.57 | 0.45 | 0.48 | 1.20 | 0.81 |
| 2adf | 6.20 | 0.39 | 0.78 | 0.56 | 2.11 | 0.39 | 0.42 | 0.91 | 0.43 |
| 2d7t | 13.32 | 0.82 | 1.07 | 0.87 | 5.90 | 0.53 | 0.56 | 0.79 | 0.98 |
| 2e27 | 4.19 | 0.33 | 0.47 | 0.36 | 3.03 | 0.57 | 0.30 | 0.63 | 1.46 |
| 2fb4 | 2.54 | 0.42 | 1.17 | 0.58 | 8.25 | 1.15 | 1.26 | 0.78 | 0.64 |
| 2fbj | 12.86 | 0.48 | 0.83 | 0.57 | 1.38 | 0.48 | 1.15 | 0.89 | 2.18 |
| 2r8s | 6.83 | 0.62 | 1.93 | 1.76 | 2.76 | 0.52 | 3.04 | 0.38 | 0.57 |
| 2v17 | 9.30 | 0.58 | 0.86 | 1.60 | 1.72 | 0.68 | 0.63 | 0.63 | 0.63 |
| 2vxv | 0.86 | 0.37 | 0.90 | 1.00 | 4.40 | 0.42 | 0.30 | 0.64 | 0.62 |
| 2w60 | 2.59 | 0.54 | 1.17 | 0.62 | 0.77 | 0.74 | 0.59 | 1.25 | 0.37 |
| 2xwt | 4.50 | 0.44 | 1.41 | 0.43 | 2.48 | 0.57 | 0.74 | 0.54 | 2.08 |
| 2ypv | 5.19 | 0.71 | 1.03 | 0.84 | 6.09 | 0.48 | 0.35 | 0.84 | 0.59 |
| 3e8u | 2.87 | 0.61 | 0.84 | 0.66 | 2.32 | 0.41 | 0.67 | 0.71 | 0.67 |
| 3eo9 | 5.22 | 0.49 | 1.47 | 0.91 | 2.06 | 0.43 | 0.36 | 0.78 | 0.69 |
| 3g5y | 2.18 | 0.29 | 0.75 | 0.39 | 0.42 | 0.73 | 0.48 | 0.78 | 0.46 |
| 3giz | 2.69 | 0.48 | 0.74 | 0.55 | 4.86 | 0.48 | 1.98 | 0.79 | 0.68 |
| 3gnm | 1.96 | 0.31 | 0.62 | 0.51 | 2.45 | 0.44 | 0.66 | 0.70 | 0.79 |
| 3go1 | 6.31 | 0.47 | 1.61 | 2.44 | 6.56 | 0.38 | 0.60 | 0.82 | 2.89 |
| 3hc4 | 7.98 | 1.03 | 0.57 | 1.13 | 1.39 | 0.46 | 0.55 | 0.32 | 0.26 |
| 3hnt | 0.36 | 0.62 | 1.24 | 0.74 | 3.27 | 0.63 | 0.43 | 1.05 | 0.90 |
| 3i9g | 5.75 | 0.52 | 0.44 | 1.06 | 2.93 | 0.61 | 0.75 | 0.78 | 0.80 |
| 3liz | 2.71 | 0.58 | 1.41 | 0.34 | 4.57 | 0.75 | 0.34 | 0.58 | 0.37 |
| 3lmj | 0.75 | 0.73 | 3.21 | 6.59 | 6.60 | 0.48 | 0.58 | 0.77 | 1.71 |
| 3m8o | 8.76 | 0.64 | 2.22 | 1.04 | 8.38 | 0.46 | 1.27 | 1.03 | 1.10 |
| 3mlr | 3.72 | 0.57 | 0.70 | 1.12 | 3.04 | 5.30 | 3.16 | 1.63 | - |
| 3mxw | 2.71 | 1.21 | 1.31 | 1.29 | 1.72 | 0.41 | 0.74 | 0.66 | 0.80 |
| 3nps | 3.59 | 1.27 | 3.74 | 2.99 | 5.53 | 0.61 | 1.02 | 0.87 | 2.64 |
| 3oz9 | 10.17 | 0.54 | 0.79 | 0.88 | 2.16 | 0.61 | 0.98 | 0.94 | 2.15 |
| 3p0y | 3.34 | 0.38 | 2.74 | 2.61 | 2.60 | 0.54 | 0.30 | 0.71 | 1.77 |
| 3t65 | 1.81 | 0.35 | 0.93 | 0.75 | 0.44 | 0.41 | 0.67 | 0.50 | 0.69 |
| 3umt | 7.46 | 0.59 | 1.59 | 0.97 | 4.04 | 0.55 | 0.50 | 0.67 | 0.83 |
| 3v0w | 7.14 | 0.56 | 1.53 | 1.59 | 1.74 | 0.41 | 0.51 | 0.63 | 0.99 |
| 4f57 | 4.10 | 0.58 | 1.17 | 2.46 | 8.28 | 0.69 | 1.01 | 0.92 | 1.41 |
| 4h0h | 5.66 | 0.52 | 0.65 | 0.67 | 2.67 | 0.58 | 0.61 | 0.76 | 0.74 |
| 4h20 | 3.67 | 0.54 | 0.86 | 0.35 | 1.75 | 0.58 | 0.50 | 0.86 | 1.57 |
| 4hpy | 3.78 | 0.38 | 2.77 | 0.61 | 0.61 | 0.54 | 0.75 | 0.97 | 1.18 |
| 4nzu | 5.55 | 0.47 | 0.83 | 0.77 | 7.16 | 0.71 | 0.45 | 1.12 | 0.82 |
| **Mean** | 5.19 | 0.57 | 1.22 | 1.14 | 3.48 | 0.67 | 0.80 | 0.87 | 1.06 |
| **SD** | 2.96 | 0.20 | 0.71 | 1.01 | 2.21 | 0.71 | 0.63 | 0.34 | 0.61 |

**Table S4. CDR loop RMSD and OCD results for RosettaAntibody-G on therapeutic benchmark.**

| Target | OCD | H Fr (Å) | H1 (Å) | H2 (Å) | H3 (Å) | L Fr (Å) | L1 (Å) | L2 (Å) | L3 (Å) |
|---|---|---|---|---|---|---|---|---|---|
| 1bey | 4.23 | 0.96 | 1.81 | 2.06 | 3.58 | 0.67 | 0.72 | 1.20 | 1.22 |
| 1cz8 | 3.32 | 0.35 | 2.82 | 0.30 | 0.40 | 0.45 | 0.75 | 0.96 | 0.61 |
| 1mim | 13.64 | 0.63 | 1.00 | 0.74 | 0.85 | 0.63 | 1.35 | 0.98 | 2.36 |
| 1sy6 | 3.53 | 0.32 | 0.88 | 0.46 | 1.62 | 0.49 | 0.52 | 0.80 | 0.84 |
| 1yy8 | 5.26 | 1.17 | 0.99 | 1.04 | 4.84 | 0.55 | 0.58 | 0.77 | 0.71 |
| 2hwz | 1.74 | 0.31 | 1.05 | 0.42 | 1.79 | 0.56 | 0.58 | 1.06 | 2.23 |
| 3eo0 | 5.58 | 0.72 | 2.81 | 1.61 | 9.58 | 0.39 | 1.57 | 1.13 | 0.83 |
| 3gkw | 10.31 | 1.23 | 2.68 | 0.94 | 7.81 | 0.65 | 1.53 | 1.15 | 0.94 |
| 3nfs | 4.31 | 0.84 | 1.80 | 1.65 | 3.34 | 0.46 | 1.00 | 0.91 | 1.38 |
| 3o2d | 4.18 | 0.56 | 0.86 | 0.94 | 4.71 | 0.55 | 0.82 | 0.83 | 0.40 |
| 3pp3 | 2.22 | 0.53 | 0.90 | 1.30 | 4.15 | 0.54 | 1.06 | 0.90 | 0.92 |
| 3qwo | 4.57 | 0.35 | 0.66 | 1.32 | 1.73 | 0.56 | 0.71 | 1.18 | 2.22 |
| 3u0t | 17.22 | 0.66 | 2.22 | 1.15 | 5.15 | 0.71 | 1.17 | 0.96 | 1.40 |
| 4cni | 9.41 | 0.58 | 0.64 | 1.30 | 3.42 | 0.52 | 0.34 | 0.79 | 0.81 |
| 4dn3 | 7.94 | 0.68 | 3.74 | 1.30 | 2.66 | 0.41 | 1.44 | 0.89 | 1.02 |
| 4g5z | 4.56 | 0.38 | 0.79 | 0.40 | 0.56 | 0.50 | 0.31 | 0.73 | 0.53 |
| 4g6k | 4.11 | 0.38 | 0.76 | 1.24 | 2.33 | 0.59 | 0.73 | 0.69 | 0.53 |
| 4hkz | 4.75 | 1.07 | 1.01 | 1.00 | 3.55 | 0.59 | 0.43 | 0.46 | 0.65 |
| 4i77 | 4.27 | 0.87 | 0.90 | 1.09 | 3.10 | 0.35 | 1.10 | 0.66 | 0.53 |
| 4irz | 2.62 | 0.64 | 1.02 | 1.26 | 3.33 | 0.58 | 0.70 | 0.80 | 1.30 |
| 4kaq | 3.65 | 1.02 | 1.12 | 0.79 | 2.82 | 0.48 | 0.36 | 0.61 | 0.94 |
| 4m6n | 4.57 | 0.31 | 0.66 | 0.66 | 3.81 | 0.66 | 1.27 | 0.88 | 1.57 |
| 4nyl | 3.47 | 0.54 | 1.02 | 0.76 | 4.36 | 0.77 | 0.98 | 0.77 | 0.82 |
| 4od2 | 5.18 | 0.69 | 0.88 | 0.60 | 8.60 | 0.63 | 0.74 | 0.96 | 1.72 |
| 4ojf | 4.80 | 0.45 | 1.05 | 1.22 | 4.98 | 0.48 | 0.27 | 1.12 | 0.73 |
| 4qxg | 3.26 | 0.54 | 0.97 | 0.79 | 1.76 | 0.44 | 0.39 | 0.48 | 2.14 |
| 4x7s | 5.23 | 0.62 | 1.52 | 3.26 | 2.46 | 0.57 | 0.60 | 0.60 | 0.60 |
| 4ypg | 2.84 | 0.47 | 1.18 | 1.20 | 4.51 | 0.34 | 0.54 | 0.63 | 1.23 |
| 5csz | 5.96 | 0.59 | 0.71 | 1.32 | 2.69 | 0.39 | 0.75 | 0.38 | 1.31 |
| 5dk3 | 1.27 | 0.56 | 2.54 | 0.73 | 0.94 | 0.34 | 0.85 | 0.52 | 0.33 |
| 5ggq | 1.15 | 0.46 | 0.89 | 0.75 | 2.17 | 0.46 | 1.93 | 0.89 | 0.38 |
| 5ggu | 6.62 | 0.97 | 1.20 | 1.00 | 8.20 | 0.43 | 0.43 | 0.81 | 0.61 |
| 5i5k | 8.32 | 1.10 | 0.92 | 0.96 | 5.21 | 0.89 | 0.99 | 0.74 | 1.57 |
| 5jxe | 3.75 | 0.50 | 3.68 | 1.66 | 5.74 | 0.68 | 1.23 | 0.68 | 0.73 |
| 5kmv | 11.64 | 0.60 | 1.57 | 0.87 | 1.28 | 0.48 | 0.86 | 1.07 | 0.53 |
| 5l6y | 11.90 | 0.36 | 0.72 | 0.97 | 12.57 | 0.50 | 0.58 | 0.82 | 1.87 |
| 5n2k | 4.14 | 0.47 | 0.78 | 0.49 | 3.91 | 0.62 | 1.08 | 1.01 | 6.54 |
| 5nhw | 3.48 | 0.46 | 2.54 | 0.68 | 1.24 | 0.68 | 0.81 | 0.82 | 5.18 |
| 5sx4 | 7.47 | 1.09 | 1.32 | 1.03 | 1.42 | 1.13 | 1.58 | 0.89 | 10.26 |
| 5tru | 4.89 | 0.46 | 0.70 | 0.51 | 3.24 | 0.36 | 1.84 | 0.59 | 1.55 |
| 5vh3 | 2.71 | 0.51 | 1.41 | 0.70 | 3.10 | 0.56 | 0.70 | 0.76 | 0.40 |
| 5wuv | 2.54 | 0.42 | 0.90 | 0.74 | 3.70 | 0.38 | 0.46 | 0.48 | 1.19 |
| 5xxy | 3.82 | 0.55 | 1.46 | 2.02 | 2.63 | 0.55 | 1.19 | 0.49 | 0.53 |
| 5y9k | 7.84 | 0.76 | 2.97 | 1.20 | 7.23 | 0.70 | 1.12 | 1.68 | 1.63 |
| 6and | 6.16 | 0.44 | 1.62 | 0.79 | 2.69 | 0.53 | 1.07 | 0.91 | 0.67 |
| **Mean** | 5.43 | 0.63 | 1.42 | 1.05 | 3.77 | 0.55 | 0.89 | 0.83 | 1.48 |
| **SD** | 3.32 | 0.25 | 0.83 | 0.52 | 2.53 | 0.15 | 0.41 | 0.24 | 1.76 |

**Table S5. CDR loop RMSD and OCD results for RepertoireBuilder on RosettaAntibody benchmark.**

| Target | OCD | H Fr (Å) | H1 (Å) | H2 (Å) | H3 (Å) | L Fr (Å) | L1 (Å) | L2 (Å) | L3 (Å) |
|---|---|---|---|---|---|---|---|---|---|
| 1dlf | 10.60 | 0.86 | 0.84 | 1.02 | 2.93 | 0.60 | 0.39 | 0.51 | 0.40 |
| 1fns | 5.37 | 0.51 | 0.54 | 1.45 | 3.99 | 0.55 | 0.53 | 0.24 | 0.42 |
| 1gig | 5.07 | 0.60 | 0.58 | 0.85 | 3.11 | 0.40 | 0.59 | 1.09 | 0.93 |
| 1jfq | 3.60 | 0.65 | 0.51 | 1.15 | 1.88 | 0.34 | 0.36 | 0.41 | 0.28 |
| 1jpt | 1.47 | 0.51 | 0.37 | 0.72 | 0.53 | 0.40 | 0.59 | 0.55 | 0.38 |
| 1mfa | 7.63 | 0.61 | 0.52 | 0.82 | 0.41 | 0.45 | 0.70 | 0.56 | 0.49 |
| 1mlb | 4.94 | 0.58 | 0.52 | 1.68 | 0.65 | 0.57 | 0.56 | 0.90 | 0.60 |
| 1mqk | 4.48 | 0.34 | 0.53 | 0.83 | 1.92 | 0.46 | 0.55 | 0.50 | 2.04 |
| 1nlb | 9.29 | 0.46 | 0.72 | 0.50 | 1.79 | 0.66 | 0.44 | 0.49 | 0.72 |
| 1oaq | 5.63 | 0.55 | 0.50 | 0.58 | 2.14 | 0.50 | 0.58 | 0.63 | 1.02 |
| 1seq | 7.46 | 0.62 | 0.65 | 0.72 | 3.71 | 0.37 | 0.66 | 0.35 | 0.86 |
| 2adf | 4.67 | 0.31 | 0.60 | 0.86 | 2.51 | 0.34 | 0.46 | 0.49 | 0.32 |
| 2d7t | 8.09 | 0.71 | 1.13 | 0.65 | 3.73 | 0.54 | 0.76 | 0.42 | 1.06 |
| 2e27 | 8.15 | 0.66 | 0.63 | 0.51 | 4.62 | 0.48 | 0.36 | 0.52 | 2.40 |
| 2fb4 | 1.45 | 0.43 | 0.43 | 0.64 | 5.29 | 0.98 | 0.79 | 0.50 | 1.45 |
| 2fbj | 7.51 | 0.51 | 0.94 | 0.87 | 2.37 | 0.38 | 1.16 | 0.87 | 2.10 |
| 2r8s | 1.50 | 0.50 | 0.88 | 1.86 | 2.50 | 0.45 | 0.88 | 0.62 | 1.04 |
| 2v17 | 9.98 | 0.42 | 0.50 | 1.20 | 2.34 | 0.78 | 1.03 | 0.62 | 0.65 |
| 2vxv | 1.91 | 0.22 | 0.72 | 0.74 | 3.92 | 0.53 | 0.39 | 0.92 | 2.62 |
| 2w60 | 4.14 | 0.48 | 0.74 | 0.50 | 2.46 | 0.42 | 0.69 | 0.25 | 0.63 |
| 2xwt | 8.67 | 0.39 | 1.43 | 0.44 | 3.63 | 0.53 | 0.36 | 0.87 | 1.51 |
| 2ypv | 6.75 | 1.04 | 0.74 | 0.92 | 3.88 | 0.69 | 0.91 | 0.39 | 0.70 |
| 3e8u | 4.91 | 0.73 | 0.50 | 0.66 | 1.54 | 0.42 | 0.65 | 0.45 | 0.68 |
| 3eo9 | 5.64 | 0.39 | 0.87 | 0.58 | 3.39 | 0.48 | 0.50 | 0.52 | 0.70 |
| 3g5y | 7.72 | 0.60 | 0.48 | 0.36 | 1.04 | 0.67 | 0.48 | 0.43 | 0.47 |
| 3giz | 4.82 | 0.67 | 0.80 | 0.56 | 2.58 | 0.48 | 0.47 | 0.57 | 0.55 |
| 3gnm | 3.65 | 0.76 | 0.58 | 0.99 | 1.41 | 0.47 | 0.70 | 0.27 | 0.71 |
| 3go1 | 3.17 | 0.43 | 2.23 | 0.95 | 3.48 | 0.65 | 1.05 | 0.95 | 3.52 |
| 3hc4 | 3.15 | 1.05 | 0.46 | 1.13 | 0.92 | 0.53 | 0.52 | 0.37 | 0.20 |
| 3hnt | 4.28 | 0.59 | 0.65 | 0.84 | 1.68 | 0.59 | 0.40 | 0.93 | 1.87 |
| 3i9g | 2.80 | 0.29 | 0.50 | 0.55 | 3.85 | 0.57 | 0.64 | 0.46 | 1.00 |
| 3liz | 5.59 | 0.29 | 0.73 | 0.50 | 4.43 | 0.69 | 0.33 | 0.36 | 0.40 |
| 3lmj | 2.61 | 0.99 | 3.21 | 7.12 | 6.56 | 0.45 | 0.49 | 0.43 | 2.40 |
| 3m8o | 11.24 | 0.65 | 1.92 | 1.15 | 7.45 | 0.79 | 1.64 | 0.44 | 0.67 |
| 3mlr | 5.69 | 0.52 | 0.25 | 1.27 | 5.60 | 0.59 | 0.82 | 0.47 | - |
| 3mxw | 4.49 | 0.92 | 0.80 | 1.30 | 1.94 | 0.30 | 0.39 | 0.38 | 0.66 |
| 3nps | 4.87 | 0.73 | 2.83 | 0.92 | 4.31 | 0.68 | 1.24 | 0.50 | 1.45 |
| 3oz9 | 4.49 | 0.62 | 0.52 | 1.70 | 2.98 | 0.54 | 0.62 | 0.24 | 1.76 |
| 3p0y | 6.18 | 0.52 | 1.15 | 1.60 | 3.11 | 0.38 | 0.58 | 0.25 | 1.40 |
| 3t65 | 3.02 | 0.42 | 0.27 | 0.66 | 0.43 | 0.36 | 0.75 | 1.04 | 0.40 |
| 3umt | 8.31 | 0.74 | 1.22 | 0.59 | 3.47 | 0.44 | 0.52 | 0.27 | 0.42 |
| 3v0w | 4.86 | 0.49 | 0.80 | 0.67 | 2.12 | 0.46 | 0.60 | 0.51 | 0.79 |
| 4f57 | 3.20 | 0.28 | 0.45 | 0.76 | 0.57 | 0.22 | 0.44 | 0.21 | 0.50 |
| 4h0h | 5.21 | 0.64 | 0.51 | 0.68 | 3.01 | 0.62 | 0.36 | 0.38 | 0.36 |
| 4h20 | 3.30 | 0.79 | 0.51 | 0.50 | 4.45 | 0.33 | 0.67 | 0.73 | 1.51 |
| 4hpy | 2.38 | 0.64 | 2.61 | 0.56 | 2.45 | 0.30 | 0.50 | 0.35 | 1.42 |
| 4nzu | 3.24 | 0.59 | 0.65 | 1.03 | 5.11 | 0.49 | 0.49 | 0.41 | 0.91 |
| **Mean** | 5.26 | 0.58 | 0.86 | 1.00 | 2.94 | 0.51 | 0.63 | 0.52 | 1.03 |
| **SD** | 2.46 | 0.20 | 0.65 | 0.98 | 1.59 | 0.15 | 0.26 | 0.23 | 0.73 |

**Table S6. CDR loop RMSD and OCD results for RepertoireBuilder on therapeutic benchmark.**

| Target | OCD | H Fr (Å) | H1 (Å) | H2 (Å) | H3 (Å) | L Fr (Å) | L1 (Å) | L2 (Å) | L3 (Å) |
|---|---|---|---|---|---|---|---|---|---|
| 1bey | 2.80 | 0.54 | 1.49 | 0.91 | 3.96 | 0.68 | 0.91 | 0.70 | 0.61 |
| 1cz8 | 0.59 | 0.26 | 0.82 | 0.75 | 3.66 | 0.39 | 0.35 | 0.36 | 0.42 |
| 1mim | 3.42 | 0.56 | 0.64 | 0.92 | 0.63 | 0.38 | 0.83 | 0.62 | 1.14 |
| 1sy6 | 3.71 | 0.71 | 0.99 | 1.04 | 3.17 | 0.41 | 0.26 | 0.48 | 0.37 |
| 1yy8 | 6.38 | 0.61 | 0.54 | 0.55 | 4.21 | 0.35 | 0.45 | 0.52 | 0.91 |
| 2hwz | 4.98 | 0.28 | 0.89 | 1.31 | 2.33 | 0.44 | 0.90 | 0.59 | 2.27 |
| 3eo0 | 3.96 | 0.34 | 3.13 | 1.15 | 4.38 | 0.30 | 1.43 | 0.30 | 0.74 |
| 3gkw | 15.79 | 1.40 | 2.74 | 1.62 | 9.66 | 0.56 | 1.64 | 0.67 | 0.60 |
| 3nfs | 2.53 | 0.57 | 0.60 | 0.69 | 1.91 | 0.54 | 0.56 | 0.61 | 0.60 |
| 3o2d | 10.03 | 0.93 | 0.71 | 0.97 | 2.75 | 0.46 | 0.47 | 0.30 | 0.43 |
| 3pp3 | 4.02 | 0.29 | 0.46 | 1.17 | 3.36 | 0.64 | 0.62 | 0.59 | 0.81 |
| 3qwo | 3.14 | 0.36 | 0.34 | 1.34 | 2.77 | 0.38 | 0.76 | 0.54 | 0.66 |
| 3u0t | 10.10 | 0.67 | 1.62 | 0.91 | 2.29 | 0.50 | 1.05 | 0.47 | 0.93 |
| 4cni | 1.46 | 0.60 | 0.89 | 1.26 | 1.28 | 0.63 | 0.51 | 0.52 | 0.59 |
| 4dn3 | 2.36 | 0.49 | 1.54 | 0.93 | 1.74 | 0.33 | 1.36 | 0.52 | 1.44 |
| 4g5z | 6.55 | 0.49 | 0.84 | 0.57 | 1.51 | 0.61 | 0.40 | 0.69 | 0.68 |
| 4g6k | 2.36 | 0.62 | 0.51 | 1.22 | 2.95 | 0.67 | 0.57 | 0.46 | 0.93 |
| 4hkz | 2.29 | 0.26 | 0.42 | 0.83 | 2.98 | 0.37 | 0.38 | 0.48 | 0.61 |
| 4i77 | 5.19 | 0.29 | 1.22 | 0.87 | 3.39 | 0.40 | 0.49 | 0.48 | 0.68 |
| 4irz | 3.99 | 0.99 | 1.26 | 0.60 | 3.98 | 0.46 | 0.71 | 0.40 | 1.55 |
| 4kaq | 4.21 | 0.75 | 0.47 | 0.89 | 2.79 | 0.70 | 0.80 | 0.57 | 0.80 |
| 4m6n | 4.11 | 0.31 | 0.76 | 0.36 | 2.51 | 0.34 | 1.88 | 0.35 | 1.65 |
| 4nyl | 7.73 | 0.74 | 0.45 | 0.67 | 5.12 | 0.71 | 0.63 | 0.69 | 1.05 |
| 4od2 | 0.89 | 0.63 | 0.49 | 0.76 | 6.49 | 0.51 | 1.11 | 1.25 | 2.14 |
| 4ojf | 6.62 | 0.76 | 0.59 | 1.32 | 3.84 | 0.39 | 0.58 | 0.40 | 0.57 |
| 4qxg | 0.82 | 0.42 | 0.45 | 0.76 | 2.06 | 0.47 | 0.52 | 0.66 | 1.81 |
| 4x7s | 6.16 | 0.77 | 1.72 | 1.18 | 2.71 | 0.45 | 0.45 | 0.28 | 0.69 |
| 4ypg | 4.22 | 1.05 | 1.15 | 1.09 | 2.87 | 0.49 | 0.33 | 0.93 | 0.82 |
| 5csz | 2.55 | 0.56 | 0.74 | 0.66 | 2.67 | 0.26 | 0.50 | 0.33 | 1.43 |
| 5dk3 | 5.32 | 0.80 | 0.61 | 1.20 | 2.80 | 0.49 | 0.46 | 0.70 | 0.40 |
| 5ggq | 1.88 | 0.64 | 0.47 | 0.73 | 0.55 | 0.51 | 0.36 | 0.63 | 0.51 |
| 5ggu | 4.40 | 0.38 | 0.52 | 0.48 | 6.34 | 0.51 | 0.49 | 0.35 | 0.92 |
| 5i5k | 3.29 | 1.16 | 0.62 | 1.03 | 4.87 | 0.90 | 0.87 | 0.61 | 1.10 |
| 5jxe | 3.76 | 0.79 | 0.57 | 1.50 | 2.86 | 0.66 | 0.54 | 0.61 | 0.62 |
| 5kmv | 3.52 | 0.35 | 0.66 | 0.54 | 0.56 | 0.22 | 0.44 | 0.37 | 0.40 |
| 5l6y | 1.88 | 0.96 | 0.91 | 0.63 | 5.61 | 0.32 | 0.31 | 0.53 | 1.99 |
| 5n2k | 1.65 | 0.62 | 0.94 | 0.88 | 3.28 | 0.48 | 0.69 | 0.21 | 4.27 |
| 5nhw | 3.38 | 0.99 | 0.66 | 0.84 | 1.22 | 0.27 | 1.08 | 0.21 | 4.76 |
| 5sx4 | 7.62 | 1.29 | 0.94 | 1.40 | 2.49 | 0.46 | 0.61 | 0.62 | 0.79 |
| 5tru | 1.95 | 0.36 | 0.32 | 0.98 | 2.32 | 0.33 | 1.72 | 0.33 | 0.96 |
| 5vh3 | 4.22 | 0.27 | 0.60 | 0.63 | 1.62 | 0.56 | 0.48 | 0.58 | 0.70 |
| 5wuv | 1.98 | 0.63 | 0.88 | 0.78 | 3.00 | 0.38 | 0.23 | 0.34 | 0.41 |
| 5xxy | 4.89 | 0.51 | 0.66 | 2.10 | 2.37 | 0.46 | 1.07 | 0.62 | 0.43 |
| 5y9k | 7.48 | 0.59 | 2.79 | 0.74 | 4.13 | 0.34 | 0.32 | 0.35 | 1.90 |
| 6and | 6.29 | 0.45 | 0.52 | 1.48 | 2.80 | 0.47 | 0.95 | 0.37 | 0.41 |
| **Mean** | 4.37 | 0.62 | 0.91 | 0.96 | 3.13 | 0.47 | 0.71 | 0.52 | 1.08 |
| **SD** | 2.85 | 0.28 | 0.63 | 0.35 | 1.68 | 0.14 | 0.40 | 0.19 | 0.90 |

**Table S7. CDR loop RMSD and OCD results for ABodyBuilder on RosettaAntibody benchmark.**

| Target | OCD | H Fr (Å) | H1 (Å) | H2 (Å) | H3 (Å) | L Fr (Å) | L1 (Å) | L2 (Å) | L3 (Å) |
|---|---|---|---|---|---|---|---|---|---|
| 1dlf | 8.05 | 0.66 | 0.93 | 0.47 | 4.19 | 0.63 | 1.10 | 0.49 | 0.80 |
| 1fns | 3.07 | 0.38 | 0.52 | 0.52 | 5.44 | 0.55 | 0.83 | 0.42 | 0.84 |
| 1gig | 11.80 | 0.54 | 0.99 | 0.62 | 4.02 | 0.44 | 0.55 | 1.09 | 0.54 |
| 1jfq | 5.43 | 0.64 | 0.53 | 1.17 | 1.25 | 0.49 | 0.24 | 0.23 | 0.27 |
| 1jpt | 3.00 | 0.60 | 0.76 | 1.32 | 1.28 | 0.40 | 1.05 | 0.50 | 0.69 |
| 1mfa | 2.28 | 0.42 | 0.38 | 0.38 | 0.17 | 0.23 | 0.70 | 0.23 | 0.37 |
| 1mlb | 4.00 | 0.57 | 0.72 | 1.01 | 0.84 | 0.41 | 0.47 | 0.65 | 0.56 |
| 1mqk | 4.24 | 0.37 | 0.43 | 0.44 | 2.37 | 0.42 | 1.02 | 0.42 | 2.13 |
| 1nlb | 3.54 | 0.31 | 0.50 | 0.31 | 2.42 | 0.42 | 1.05 | 0.21 | 0.88 |
| 1oaq | 3.29 | 0.62 | 0.54 | 0.66 | 1.83 | 0.42 | 0.55 | 0.51 | 1.27 |
| 1seq | 9.24 | 0.33 | 0.55 | 1.02 | 5.12 | 0.32 | 0.80 | 0.26 | 1.17 |
| 2adf | 4.38 | 0.21 | 0.73 | 0.74 | 2.07 | 0.54 | 0.60 | 0.33 | 0.53 |
| 2d7t | 13.72 | 0.76 | 0.69 | 0.82 | 2.88 | 0.48 | 0.73 | 0.37 | 1.26 |
| 2e27 | 4.52 | 0.31 | 0.49 | 0.42 | 4.56 | 0.64 | 0.38 | 0.61 | 2.37 |
| 2fb4 | 2.92 | 0.51 | 0.50 | 0.46 | 5.63 | 0.77 | 0.92 | 0.49 | 1.31 |
| 2fbj | 6.53 | 0.40 | 0.43 | 0.53 | 1.99 | 0.49 | 1.18 | 0.92 | 2.24 |
| 2r8s | 1.63 | 0.70 | 4.01 | 2.25 | 6.93 | 0.45 | 0.42 | 0.62 | 0.58 |
| 2v17 | 3.25 | 0.46 | 0.49 | 1.20 | 2.66 | 0.72 | 0.89 | 0.60 | 0.91 |
| 2vxv | 1.52 | 0.25 | 0.79 | 0.71 | 4.68 | 0.54 | 0.59 | 0.91 | 3.48 |
| 2w60 | 10.18 | 0.30 | 0.46 | 0.65 | 2.49 | 0.60 | 0.57 | 0.66 | 0.35 |
| 2xwt | 8.91 | 0.54 | 1.21 | 0.60 | 3.51 | 0.34 | 0.40 | 0.90 | 0.82 |
| 2ypv | 5.96 | 0.51 | 0.86 | 0.90 | 2.67 | 0.61 | 0.33 | 0.28 | 0.78 |
| 3e8u | 1.81 | 0.56 | 0.58 | 0.98 | 2.24 | 0.31 | 0.53 | 0.50 | 0.40 |
| 3eo9 | 5.51 | 0.42 | 0.57 | 1.51 | 3.04 | 0.54 | 0.57 | 0.59 | 1.49 |
| 3g5y | 4.70 | 0.24 | 0.39 | 0.38 | 0.94 | 0.71 | 0.28 | 0.49 | 0.60 |
| 3giz | 2.92 | 0.40 | 0.38 | 0.51 | 2.94 | 0.52 | 0.55 | 0.53 | 0.56 |
| 3gnm | 5.54 | 0.57 | 0.69 | 0.46 | 2.17 | 0.33 | 0.67 | 0.24 | 0.75 |
| 3go1 | 6.19 | 0.53 | 1.45 | 0.87 | 3.23 | 0.65 | 1.74 | 0.90 | 3.59 |
| 3hc4 | 5.45 | 1.13 | 2.54 | 1.38 | 3.59 | 0.32 | 0.53 | 0.35 | 0.65 |
| 3hnt | 2.20 | 0.58 | 0.29 | 0.49 | 2.27 | 0.56 | 0.69 | 1.09 | 1.29 |
| 3i9g | 1.47 | 0.25 | 0.58 | 0.90 | 3.32 | 0.38 | 0.83 | 0.44 | 0.55 |
| 3liz | 2.68 | 0.48 | 0.65 | 0.54 | 4.38 | 0.61 | 0.37 | 0.27 | 0.72 |
| 3lmj | 4.17 | 0.50 | 3.10 | 2.78 | 4.88 | 0.60 | 0.59 | 0.81 | 1.02 |
| 3m8o | 3.18 | 0.93 | 2.31 | 0.86 | 5.67 | 0.74 | 3.00 | 0.56 | 2.19 |
| 3mlr | 3.98 | 0.51 | 0.80 | 1.24 | 4.93 | 0.54 | 0.75 | 0.74 | - |
| 3mxw | 2.70 | 0.92 | 2.07 | 1.26 | 2.74 | 0.57 | 0.61 | 0.55 | 0.97 |
| 3nps | 2.86 | 0.86 | 3.35 | 1.71 | 5.10 | 0.58 | 1.01 | 0.45 | 1.77 |
| 3oz9 | 6.12 | 0.49 | 0.63 | 0.55 | 2.63 | 0.68 | 0.75 | 0.69 | 2.52 |
| 3p0y | 4.34 | 0.40 | 0.91 | 1.76 | 2.57 | 0.35 | 0.46 | 0.22 | 1.58 |
| 3t65 | 1.47 | 0.23 | 0.26 | 0.32 | 0.26 | 0.37 | 0.50 | 0.41 | 0.33 |
| 3umt | 4.64 | 0.44 | 0.82 | 0.92 | 2.67 | 0.47 | 0.95 | 0.51 | 0.43 |
| 3v0w | 5.99 | 0.43 | 0.83 | 0.83 | 1.53 | 0.26 | 0.36 | 0.25 | 0.38 |
| 4f57 | 3.85 | 0.28 | 0.40 | 0.78 | 0.60 | 0.22 | 0.52 | 0.21 | 0.50 |
| 4h0h | 5.76 | 0.80 | 0.50 | 1.18 | 1.47 | 0.66 | 0.64 | 0.74 | 0.46 |
| 4h20 | 5.12 | 0.64 | 1.76 | 0.42 | 2.73 | 0.52 | 0.44 | 0.38 | 0.94 |
| 4hpy | 2.87 | 0.30 | 2.80 | 0.61 | 0.62 | 0.39 | 0.55 | 0.36 | 1.28 |
| 4nzu | 3.43 | 0.27 | 0.51 | 0.72 | 2.53 | 0.46 | 0.36 | 0.42 | 0.85 |
| **Mean** | 4.69 | 0.50 | 0.99 | 0.88 | 2.94 | 0.49 | 0.72 | 0.52 | 1.09 |
| **SD** | 2.66 | 0.20 | 0.88 | 0.51 | 1.58 | 0.14 | 0.44 | 0.23 | 0.79 |

**Table S8. CDR loop RMSD and OCD results for ABodyBuilder on therapeutic benchmark.**

| Target | OCD | H Fr (Å) | H1 (Å) | H2 (Å) | H3 (Å) | L Fr (Å) | L1 (Å) | L2 (Å) | L3 (Å) |
|---|---|---|---|---|---|---|---|---|---|
| 1bey | 2.81 | 0.56 | 1.07 | 1.31 | 3.37 | 0.66 | 0.75 | 0.82 | 1.08 |
| 1cz8 | 1.09 | 0.19 | 0.26 | 0.26 | 0.33 | 0.19 | 0.36 | 0.23 | 0.27 |
| 1mim | 4.83 | 0.54 | 0.76 | 0.60 | 0.71 | 0.41 | 0.84 | 0.49 | 2.77 |
| 1sy6 | 2.03 | 0.84 | 0.76 | 0.77 | 7.12 | 0.42 | 0.41 | 0.43 | 0.44 |
| 1yy8 | 0.93 | 0.15 | 0.14 | 0.15 | 0.12 | 0.28 | 0.16 | 0.13 | 0.13 |
| 2hwz | 1.31 | 0.27 | 0.67 | 1.31 | 1.77 | 0.47 | 0.79 | 0.61 | 2.20 |
| 3eo0 | 2.15 | 0.33 | 3.51 | 1.18 | 9.54 | 0.27 | 1.45 | 0.22 | 3.46 |
| 3gkw | 20.52 | 1.40 | 3.19 | 4.76 | 7.82 | 0.72 | 1.21 | 0.39 | 0.84 |
| 3nfs | 2.89 | 0.51 | 0.58 | 0.91 | 1.52 | 0.34 | 0.75 | 0.55 | 0.91 |
| 3o2d | 3.42 | 0.39 | 0.66 | 0.55 | 3.88 | 0.49 | 0.80 | 0.38 | 0.94 |
| 3pp3 | 3.15 | 0.28 | 0.83 | 0.98 | 3.30 | 0.50 | 0.93 | 0.51 | 1.38 |
| 3qwo | 1.39 | 0.27 | 0.67 | 1.31 | 1.77 | 0.47 | 0.79 | 0.61 | 2.20 |
| 3u0t | 10.88 | 0.72 | 1.74 | 0.96 | 3.69 | 0.55 | 0.80 | 0.87 | 1.39 |
| 4cni | 2.52 | 0.36 | 0.58 | 1.08 | 1.09 | 0.51 | 0.40 | 0.45 | 0.86 |
| 4dn3 | 3.15 | 0.48 | 2.52 | 1.94 | 1.93 | 0.48 | 1.34 | 0.47 | 1.63 |
| 4g5z | 4.89 | 0.43 | 0.37 | 0.52 | 2.15 | 0.65 | 0.87 | 0.76 | 0.71 |
| 4g6k | 7.37 | 0.55 | 2.57 | 1.30 | 1.38 | 0.73 | 0.79 | 0.26 | 1.53 |
| 4hkz | 1.50 | 0.33 | 0.41 | 0.74 | 0.15 | 0.36 | 0.37 | 0.35 | 0.74 |
| 4i77 | 1.89 | 0.29 | 1.48 | 1.19 | 2.28 | 0.37 | 0.60 | 0.50 | 1.12 |
| 4irz | 0.94 | 0.57 | 2.81 | 1.06 | 5.16 | 0.55 | 1.03 | 0.48 | 1.61 |
| 4kaq | 4.46 | 0.94 | 0.33 | 0.94 | 5.19 | 0.41 | 0.50 | 0.42 | 0.67 |
| 4m6n | 10.07 | 0.31 | 0.52 | 0.54 | 3.54 | 0.62 | 1.35 | 0.42 | 2.63 |
| 4nyl | 5.47 | 0.53 | 0.61 | 0.77 | 4.14 | 0.69 | 0.60 | 0.44 | 0.68 |
| 4od2 | 2.30 | 0.53 | 0.51 | 0.67 | 5.41 | 0.58 | 1.60 | 1.51 | 2.19 |
| 4ojf | 3.17 | 0.48 | 0.46 | 2.13 | 2.70 | 0.33 | 0.88 | 0.60 | 0.60 |
| 4qxg | 3.87 | 0.41 | 0.66 | 0.71 | 1.64 | 0.46 | 1.81 | 0.77 | 1.31 |
| 4x7s | 12.37 | 0.66 | 1.63 | 1.09 | 2.94 | 0.36 | 5.18 | 0.38 | 1.28 |
| 4ypg | 2.70 | 0.42 | 0.81 | 0.64 | 4.17 | 0.40 | 3.29 | 0.86 | 0.79 |
| 5csz | 2.19 | 0.30 | 0.51 | 0.48 | 3.69 | 0.43 | 1.82 | 0.48 | 0.74 |
| 5dk3 | 3.00 | 0.37 | 0.81 | 1.08 | 2.34 | 0.34 | 1.04 | 0.54 | 0.49 |
| 5ggq | 3.80 | 0.40 | 0.43 | 0.65 | 2.40 | 0.41 | 0.33 | 0.28 | 0.92 |
| 5ggu | 2.19 | 0.27 | 0.28 | 0.49 | 4.30 | 0.30 | 0.27 | 0.23 | 0.43 |
| 5i5k | 7.23 | 1.09 | 1.62 | 2.41 | 2.17 | 0.83 | 1.02 | 0.67 | 1.07 |
| 5jxe | 3.65 | 0.35 | 0.58 | 0.84 | 2.43 | 0.63 | 0.97 | 0.64 | 0.80 |
| 5kmv | 6.28 | 0.62 | 0.89 | 1.08 | 0.71 | 0.38 | 0.77 | 0.54 | 1.30 |
| 5l6y | 3.69 | 0.28 | 0.70 | 0.62 | 3.76 | 0.27 | 0.24 | 0.28 | 2.16 |
| 5n2k | 2.24 | 0.36 | 0.20 | 0.38 | 2.52 | 0.44 | 0.44 | 0.40 | 5.24 |
| 5nhw | 3.76 | 0.27 | 0.74 | 0.61 | 0.64 | 0.27 | 0.84 | 0.21 | 4.69 |
| 5sx4 | 4.30 | 1.25 | 1.70 | 1.11 | 3.88 | 0.35 | 0.30 | 0.71 | 0.51 |
| 5tru | 5.56 | 0.34 | 0.52 | 0.47 | 2.07 | 0.28 | 1.68 | 0.41 | 1.26 |
| 5vh3 | 4.05 | 0.43 | 0.71 | 0.72 | 3.35 | 0.44 | 0.70 | 0.39 | 0.52 |
| 5wuv | 4.97 | 0.58 | 1.20 | 1.21 | 4.54 | 0.34 | 0.44 | 0.39 | 0.56 |
| 5xxy | 10.94 | 0.61 | 1.15 | 1.07 | 3.15 | 0.48 | 1.25 | 0.58 | 1.45 |
| 5y9k | 2.39 | 0.56 | 3.11 | 0.76 | 1.18 | 0.39 | 3.46 | 0.55 | 1.62 |
| 6and | 2.16 | 0.32 | 0.97 | 1.54 | 2.87 | 0.44 | 0.63 | 0.39 | 0.65 |
| **Mean** | 4.37 | 0.49 | 1.05 | 1.02 | 3.00 | 0.45 | 1.04 | 0.50 | 1.35 |
| **SD** | 3.67 | 0.26 | 0.86 | 0.73 | 1.97 | 0.14 | 0.92 | 0.23 | 1.06 |

**Appendix S1. RosettaAntibody grafting command.**

```
antibody.linuxgccrelease -fasta 1dlf.fasta -exclude_pdbs 1dlf 1c5c 2dlf 1wz1
1c5b
```
